# Nucleolar Pol II interactome reveals TBPL1, PAF1, and Pol I at intergenic rDNA drive rRNA biogenesis

**DOI:** 10.1101/2023.12.10.570972

**Authors:** Negin Khosraviani, V. Talya Yerlici, Jonathan St-Germain, Yi Yang Hou, Shi Bo Cao, Carla Ghali, Rehna Krishnan, Razqallah Hakem, Brian Raught, Karim Mekhail

## Abstract

Nucleolar ribosomal DNA (rDNA) repeats control ribosome manufacturing. rDNA harbors a ribosomal RNA (rRNA) gene and an intergenic spacer (IGS). RNA polymerase (Pol) I transcribes rRNA genes yielding the rRNA components of ribosomes. Pol II at the IGS induces rRNA production by preventing Pol I from excessively synthesizing IGS non-coding RNAs (ncRNAs) that can disrupt nucleoli. At the IGS, Pol II regulatory processes and whether Pol I function can be beneficial remain unknown. Here, we identify IGS Pol II regulators, uncovering nucleolar optimization via IGS Pol I. Compartment-enriched proximity-dependent biotin identification (compBioID) showed enrichment of the TATA-less promoter-binding TBPL1 and transcription regulator PAF1 with IGS Pol II. TBPL1 localizes to TCT motifs, driving Pol II and Pol I and maintaining its baseline ncRNA levels. PAF1 promotes Pol II elongation, preventing unscheduled R-loops that hyper-restrain IGS Pol I and its ncRNAs. PAF1 or TBPL1 deficiency disrupts nucleolar organization and rRNA biogenesis. In PAF1-deficient cells, repressing unscheduled IGS R-loops rescues nucleolar organization and rRNA production. Depleting IGS Pol I-dependent ncRNAs is sufficient to compromise nucleoli. We present the interactome of nucleolar Pol II and show its control by TBPL1 and PAF1 ensures IGS Pol I ncRNAs maintaining nucleolar structure and operation.

## Introduction

Central to ribosome biogenesis is ribosomal DNA (rDNA)^1,2,3^. In humans, ∼350 rDNA units are arranged on five chromosomes with ∼70 tandem repeats per chromosome. Each rDNA unit harbors a ∼13.3 kb ribosomal RNA (rRNA)-coding gene and a larger, ∼29.7 kb, intergenic spacer (IGS). rDNA is separated from the rest of the genome in a nuclear compartment called the nucleolus^1,4,5^. The latter is organized through the phase separation of its resident proteins into three sub-compartments—the fibrillar center (FC), dense fibrillar component (DFC), and outermost granular component (GC)^1,4,5,6,7^. At the FC-DFC boundary, RNA polymerase I (Pol I) transcribes rRNA genes into a precursor rRNA (pre-rRNA) that undergoes processing as it migrates through the DFC and GC^8,9,10^. This eventually yields the mature 18S, 5.8S, and 28S rRNAs onto which ribosomal proteins are ultimately assembled to drive universal protein synthesis and cell growth^8,9^.

We previously reported that Pol II and Pol I cohabit the IGS, where they compete to control nucleolar structure and ribosome biogenesis^11^. Operating from an IGS promoter, Pol II generates antisense intergenic non-coding RNAs (asincRNAs). These asincRNAs typically reinvade the DNA duplex behind Pol II, hybridizing with template DNA and yielding R-loop structures composed of an RNA-DNA hybrid and a single strand of DNA^11^. Pol II and its R-loops create a shield that limits Pol I recruitment to the IGS. Inhibiting Pol II hyperinduces Pol I function at the IGS^11^. Similar results were obtained upon repressing the Pol II-dependent R-loop shield at the IGS using a locus-associated R-loop repression system (LasR) employing a chimeric protein harboring dCas9 and the RNA-DNA hybrid repressor RNase H1 (RNaseH1-EGFP-dCas9 (RED))^11,12^. The unleashed Pol I synthesizes excessive levels of sense intergenic non-coding RNAs (sincRNAs) that severely disrupt nucleolar organization and ribosome biogenesis. The induction of sincRNAs preceded nucleolar disorganization, and nucleolar structure and function were rescued in IGS Pol II-deficient cells upon repressing Pol I-dependent sincRNAs using antisense oligonucleotides^11^. Thus, IGS Pol II is a key driver of ribosome biogenesis.

Chromatin immunoprecipitation (ChIP) showed that Pol II is enriched throughout the rDNA repeats peaking near the IGS 28-30 kb region^11^. ChIP sequencing (ChIP-seq) analyses also revealed that this IGS region is especially enriched in Pol II together with the intergenic promoter marks H3K27Ac, H3K9Ac, and H3K4me3. Together with strand-specific RNA analyses incorporating Pol II inhibitors, these data indicated that Pol II mostly initiates transcription from the IGS’s 28-30 kb region in the antisense orientation^11^. However, it remains unclear how IGS Pol II transcription and its impact on IGS Pol I and rRNA biogenesis are regulated. Also, whether the baseline function of IGS Pol I is biologically relevant is unknown.

Here, we reveal two nucleolar Pol II regulators, the TATA box-binding protein-like 1 (TBPL1) and Pol II-associated factor-1 (PAF1). Proteomic analyses and validations revealed these factors are enriched with Pol II at the IGS. TBPL1 deficiency restricted Pol II and Pol I recruitment and function at the IGS, reducing the levels of IGS ncRNAs. PAF1 depletion interfered with Pol II progression through the IGS, inducing unscheduled R-loops at a specific IGS region and restraining IGS Pol I function. Thus, the loss of TBPL1 or PAF1 limits IGS Pol I function. Notably, cells deficient in TBPL1 or PAF1 had altered nucleoli and rRNA biogenesis. Repression of the unscheduled IGS R-loops in PAF1-deficient cells restored IGS ncRNA expression, rescuing nucleolar organization and rRNA production. Also, directly repressing the baseline levels of IGS Pol I-dependent sincRNAs compromised nucleolar organization and rRNA biogenesis. Thus, cells have evolved mechanisms to fine-tune IGS Pol I function, the excess or deficiency of which compromises nucleolar structure and function via distinct molecular processes.

## Results

### TBPL1 and PAF1 co-enrich with nucleolar Pol II at the IGS

To uncover the nucleolar interactome of Pol II, we tagged its RPB9/POLR2I subunit with FLAG miniTurboID and used it as bait in proximity-dependent biotin identification (BioID)^13^. Sucrose gradient-based nucleolar isolation was performed and followed by mass spectrometry to identify biotinylated proteins (Fig. 1a; Supplementary Fig. 1 and 2). RPB9 was selected due to its proximity to the interface between the Pol II complex and DNA^14,15^. Non-fractionated cells served as a negative control, allowing us to identify proximity interaction candidates (PICs) of Pol II that may be enriched in nucleoli. We detected 115 PICs in the non-fractionated samples and 77 PICs in the nucleolar-enriched samples (Fig. 1b). From these 77 PICs, 55 factors were exclusively detected in the nucleolar-enriched samples. We identified several known nucleolar proteins in our nucleolar-enriched samples^16^ and known non-nucleolar proteins only in our non-nucleolar-enriched samples (Fig. 1c). STRING gene ontology analysis of the 55 PICs detected exclusively in the nucleolar-enriched samples identified 19 factors linked to nucleic acid processes (Fig. 1c, d). Further analysis of these 19 PICs showed that most of these factors are indeed involved in RNA Pol II-dependent transcription and gene expression (Fig. 1d). Amongst the 19 PICs, the polymerase-associated factor 1 (PAF1), TATA-binding protein-like 1 (TBPL1), mediator complex subunit 26 (MED26), and down-regulator of transcription 1 (DR1) were all simultaneously linked to 5/13 enriched processes and globally linked to 10/13 processes identified (Fig. 1d).

**Figure 1.**
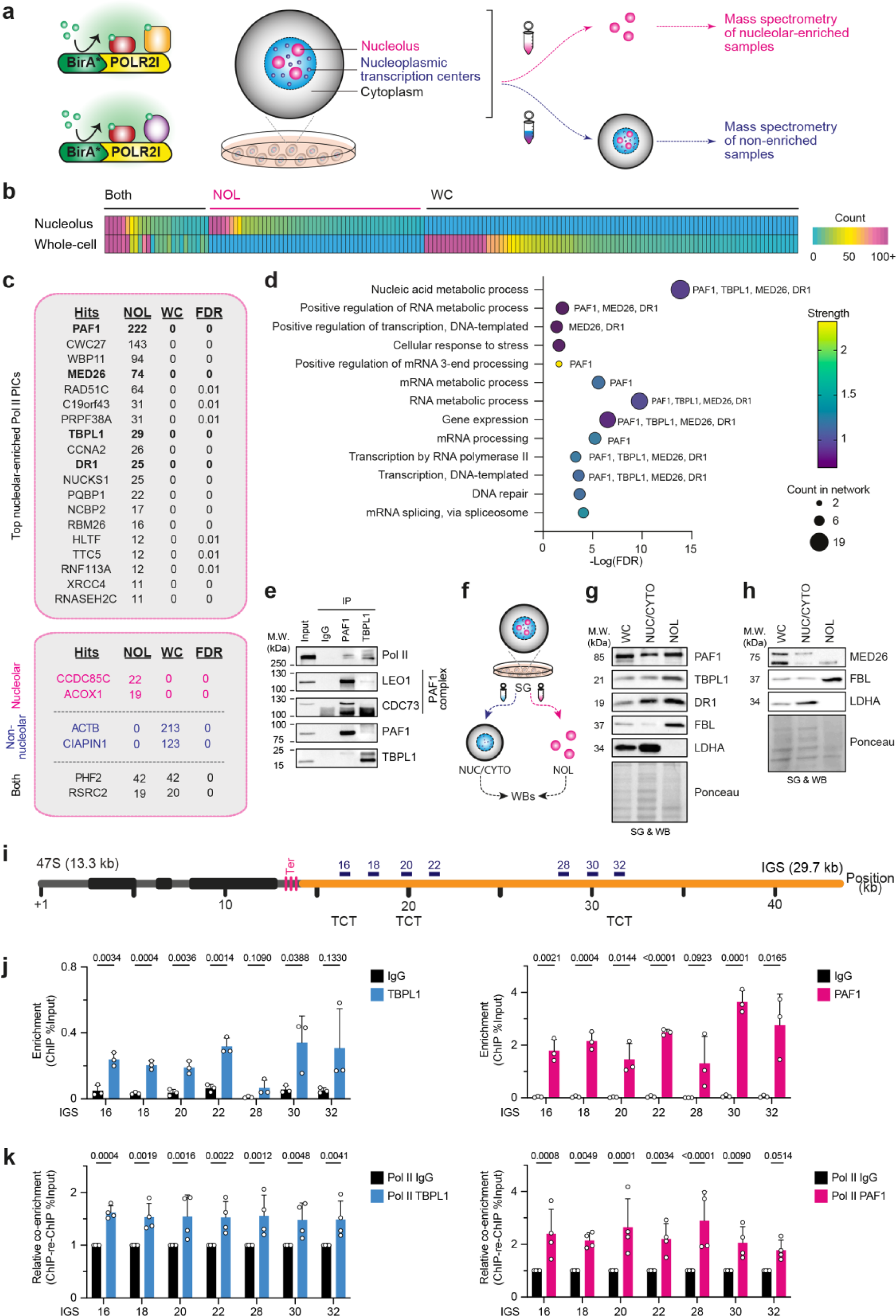
Enrichment of TBPL1 and PAF1 with nucleolar Pol II at the rDNA IGS. (**a**) Schematic of the nucleolar compBioID approach used to identify the interactome of nucleolar Pol II. Expression of the BirA-tagged Pol II POL2RI/RPB9 subunit promotes the biotinylation of proximal proteins. This is followed by sucrose gradient-based (SG) enrichment of nucleoli and use of mass spectrometry to identify biotinylated factors as proximity interaction candidates (PICs). Non-nucleolar enriched whole-cell samples served as control. (**b**) CompBioID heatmap showing peptide counts for PICs detected in nucleolar-enriched (NOL), whole-cell (WC), or both samples. (**c**) List of the top 19 nucleolar Pol II PICs detected only in the nucleolar-enriched samples and categorized under nucleic acid regulation using STRING analysis. Also shown are internal controls PICs known to be nucleolar, non-nucleolar, or both. Peptide counts for PICs detected in the NOL or WC samples are shown with the False discovery rate (FDR). (**d**) Biological processes gene ontology analysis of the 19 nucleolar-enriched Pol II PICs from panel **c**. Bubble size reflects the number of proteins in the molecular network, and colors represent the strength of the enrichment. (**e**) Immunoblots showing Pol II is pulled down with immunoprecipitated PAF1 or TBPL1. (**f**) Schematic of SG fractionation of NOL and nucleoplasmic/cytoplasmic (NUC/CYTO) fractionations followed by Western blotting (WB). (**g, h**) Immunoblots assessing the presence of PAF1, FBL, TBPL1, DR1, and MED26 in WC, NUC/CYTO, and NOL samples. Nucleolar-enriched FBL and non-nucleolar LDHA served as controls. (**i**) Schematic of a mammalian rDNA unit containing the 47S pre-rRNA-encoding 13.3 kb rRNA gene and the 29.7 kb intergenic spacer (IGS, orange) with the positioning of key primer pairs used in this study. Also shown are IGS sites harboring TCT motifs. (**j**) Chromatin immunoprecipitation (ChIP) showing the enrichment of TBPL1 (left) and PAF1 (right) at the IGS. (**k**) ChIP-re-ChIP showing the co-enrichment of TBPL1 (left) or PAF1 (right) with Pol II at the IGS. Enrichments presented relative to the Pol II+immunoglobulinG (IgG) mock ChIP-re-ChIP control. **a-k,** Experiments were performed using Flp-In 293 T-REx^TM^ (b-d) or HEK293T cells (e-k); blots (e, g, h) are representative of three biologically independent experiments; data in (j, k) are shown as mean±s.d., were from large experimental sets sharing the IgG control, were analyzed using multiple unpaired *t*-tests (j), or two-way ANOVA (k), and replicate information is *n* = 3 (j) or *n* = 4 (k) biologically independent experiments.

TBPL1 (a.k.a. TLP, TRF2, or TLF) is a TBP homolog that preferentially recognizes TCT motifs (CTCTTTCC; a.k.a. polypyrimidine initiator element)^17,18^. Recognition of TCT motifs by TBPL1 promotes Pol II transcription initiation from TATA-less promoters^17^. Once Pol II initiates transcription, the polymerase enters a stage of promoter-proximal pausing. The release of Pol II from pausing and into active elongation is regulated by PAF1, which is the major subunit of the PAF1 complex (PAF1C). The complex consists of PAF1, LEO1, CDC73, SKIC8, and CTR9, with PAF1 exhibiting the strongest interaction with Pol II^19^.

MED26 is a subunit of the mediator complex, which is recruited to promoters through direct interactions with regulatory proteins and serves as a scaffold for the assembly of a functional pre-initiation complex with Pol II and its general transcription factors^20,21,22^. DR1 (DownRegulator of transcription 1; a.k.a. Negative Cofactor 2b or NC2b), together with NC2a, forms a heterodimeric complex that binds to TBP, preventing its interaction with TFIIA or TFIIB. DR1 also binds DNA and is a component of the chromatin remodeling ATAC complex^23,24^.

Consistent with the identification of TBPL1 and PAF1 as PICs of Pol II, co-immunoprecipitation confirmed the endogenous interaction of TBPL1 and PAF1 with Pol II (Fig. 1e). The co-immunoprecipitation of LEO1 and CDC73 with PAF1 served as positive control. To validate the enrichment of TBPL1, PAF1, MED26, and DR1 in mammalian nucleoli, we performed sucrose gradient nucleolar fractionation coupled to immunoblotting (Fig. 1f). Indeed, TBPL1, PAF1, MED26, and DR1 were all present in the nucleolar fraction along with the nucleolar resident protein fibrillarin (FBL) (Fig. 1g, h). The non-nucleolar protein lactate dehydrogenase A (LDHA) was excluded from the nucleolar fraction (Fig. 1g, h). Chromatin immunoprecipitation coupled to qPCR (ChIP) revealed the enrichment of TBPL1 and PAF1 across the IGS (Fig. 1i, j), and this was comparable to other sites known to be occupied by these factors (Supplementary Fig. 3a). However, we could not detect an enrichment for MED26 or DR1 (Supplementary Fig. 3b), even though these factors were enriched at known positive control genomic sites (Supplementary Fig. 3c). To gain a better understanding of the relationship of Pol II with TBPL1 and PAF1 at the IGS, we performed sequential ChIP coupled to qPCR (ChIP-re-ChIP)^11^. In this assay, two consecutive protein pull-downs are performed before analyzing co-immunoprecipitating DNA. We found that Pol II co-enriched with TBPL1 across the IGS and with PAF1 at IGS16-30 (Fig. 1k). Further analysis of the DNA sequence at the IGS regions co-occupied by Pol II and TBPL1 identified several TCT motifs (Fig 1i, Supplementary Fig. 3d)^17,18^, further supporting the presence of TBPL1 at the IGS. These findings reveal that PAF1 and TBPL1 associate with nucleolar Pol II at the IGS.

### Dependence of intergenic Pol II and Pol I on TBPL1

To investigate the function of PAF1 and TBPL1 at the IGS, we studied TBPL1 since it has previously been reported to regulate Pol II transcription initiation from TCT-containing promoters^17,18^. We knocked down TBPL1 expression using short interfering RNA (siRNA; Fig. 2a, b). We then measured the levels of ncRNA across the IGS. Free sincRNAs are much more abundant throughout the IGS than the R-loop formation-prone asincRNAs, so most of the detectable IGS ncRNAs are Pol I-dependent transcripts^11^. Thus, to better distinguish the relationship of TBPL1 with the RNA polymerases at the IGS, we evaluated the impact of TBPL1 depletion on IGS ncRNAs in the presence or absence of Pol I or Pol II inhibition (iPol I or iPol II). To do so, cells were treated for three hours with DMSO (Veh.), low-dose actinomycin D (LAD) for iPol I, or flavopiridol (FP) for iPol II (Fig. 2c). TBPL1-depleted cells showed a significant decrease in steady-state IGS ncRNA levels, specifically at the IGS 16, 18, and 22-32 regions (Fig. 2d), partly mimicking Pol I inhibition^11^. The average change in the levels of ncRNAs at all IGS sites tested following TBPL1 knockdown was a reduction of 28.7% (Supplementary Fig. 4a). Also, iPol I was dominant over TBPL1 knockdown, almost entirely repressing ncRNA levels from IGS18 to IGS32 (Fig. 2d), as expected^11^. However, at IGS16, where iPol I did not entirely repress ncRNA levels, iPol I and TBPL1 knockdown additively lowered ncRNA levels, suggesting that these interventions decrease IGS ncRNA levels by non-identical processes (Fig. 2d). In addition, iPol II did induce ncRNA levels across the IGS as we previously showed (Fig. 2e)^11^. Notably, iPol II was fully dominant over TBPL1 depletion, except for the IGS 16 and 18 regions where TBPL1 knockdown slightly but significantly increased ncRNA levels further (Fig. 2e). Overall, these findings indicate that TBPL1 promotes IGS ncRNA expression primarily in unperturbed cells, where both Pol I and Pol II are functional at the IGS. Pol I inhibition drastically lowers ncRNA levels, muting the repressive effect of TBPL1 depletion. Similarly, in the absence of Pol II function, the drastically increased function of Pol I at the IGS greatly desensitizes the region to TBPL1 depletion.

**Figure 2.**
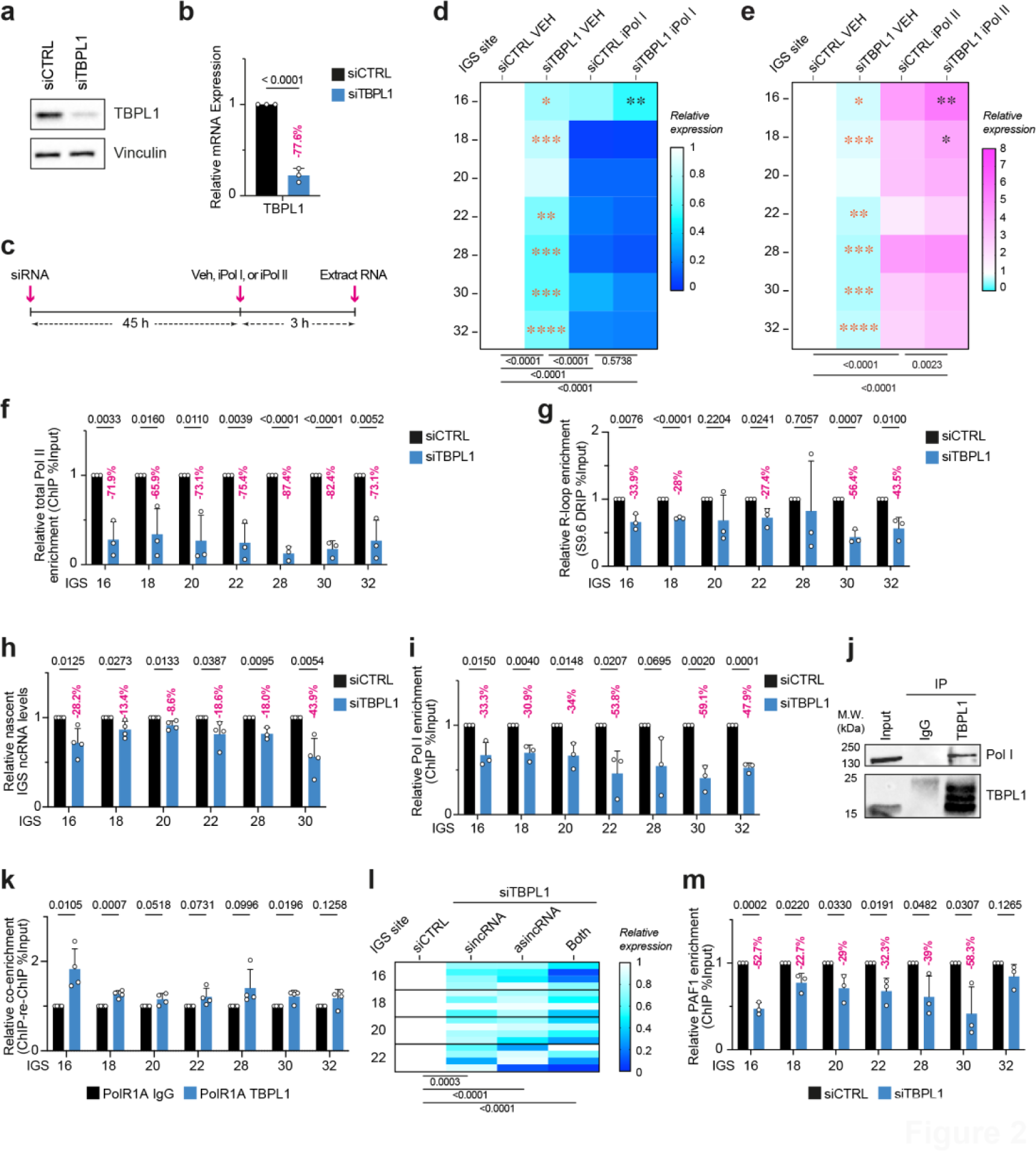
TBPL1 mediates Pol II and Pol I transcription at the rDNA IGS. (**a, b**) Validation of TBPL1 knockdown via immunoblotting (**a**) and RT-qPCR (**b**). (**c**) Schematic of the experimental setup employing a 3-hour inhibition of Pol I or Pol II (iPol I or iPol II) following siRNA-mediated knockdown of TBPL1 (siTBPL1). (**d, e**) Heatmaps showing the effect of TBPL1 knockdown on IGS ncRNA levels in cells treated with vehicle (VEH), iPol I, or iPol II. (**f**) ChIP analysis of total Pol II enrichment across the IGS. (**g**) DRIP analysis of R-loop levels at the IGS. (**h**) RT-qPCR analysis of 5-EU-labeled nascent IGS ncRNAs. (**i**) ChIP analysis of Pol I enrichment across the IGS. (**j**) Immunoblotting showing Pol I pulldown with immunoprecipitated TBPL1. (**k**) ChIP-re-ChIP assessing the co-enrichment of TBPL1 and Pol I at the IGS. (**l**) Heatmap presenting the strand-specific analysis of IGS ncRNA using ssRT-qPCR following TBPL1 knockdown. (**m**) ChIP-qPCR analysis of PAF1 enrichment at the IGS. (**a-m)** Experiments were performed using HEK293T cells; quantitative data are shown as mean±s.d. from *n* = 3 except for (l) where *n* = 4 biologically independent experiments; blots are representative of three biologically independent experiments (a, j); data in (f, i, m) were from large experimental sets sharing the IgG control; data were analyzed using unpaired *t*-tests (b, f-i, k, m), two-way ANOVA with Dunnett’s multiple comparisons tests (l), and for (d, e) shown is two-way ANOVA with Tukey’s multiple comparisons tests (below heatmaps), multiple unpaired *t*-tests comparing VEH-treated samples (orange asterisks) and two-way ANOVA comparing iPol I- or iPol II-treated samples (black asterisks).

We next sought to investigate the impact of TBPL1 on Pol I and Pol II at the IGS in unperturbed cells. TBPL1-depleted cells exhibited a significant decrease in total Pol II enrichment across the IGS (Fig. 2f). This result was similar to the impact of TBPL1 depletion on Pol II enrichment at the ribosomal protein gene *RPLP1A* (Supplementary Fig. 4b), which is transcribed via TBPL1/TCT-dependent Pol II function in *Drosophila*^25^. Decreased Pol II enrichment in TBPL1-depleted cells occurred without changing global Pol II level (Supplementary Fig. 4c). Reduced Pol II enrichment at the IGS was accompanied by a significant decrease in the levels of IGS R-loops as measured using S9.6 antibody-based DNA-RNA immunoprecipitation (DRIP; Fig. 2g; Supplementary Fig. 4d). DRIP signals were sensitive to RNaseH1 treatment *in vitro*, highlighting the specificity of R-loop detection using S9.6 (Supplementary Fig. 4d). These changes in R-loops were also observed using the GFP-tagged catalytically dead RNase H1 (GFP-dRNH1), which depicted similar R-loop levels at the IGS (Supplementary Fig. 4d-f). So, TBPL1 is required to enrich Pol II at the IGS efficiently.

Given that Pol II generates R-loops at the IGS to shield the region from excessive Pol I recruitment, we next assessed transcriptional activity at the IGS by measuring the incorporation of 5-ethynyl uridine (5-EU) in nascent transcripts^11^. TBPL1 knockdown decreased the levels of nascent IGS ncRNA (Fig. 2h, Supplementary Fig. 4g) and reduced Pol I occupancy at the IGS (Fig. 2i). This occurred without changes to global Pol I levels (Supplementary Fig. 4h). These findings point to a role for TBPL1 in enriching both Pol I and Pol II at the IGS.

Notably, co-immunoprecipitation revealed endogenous interaction between TBPL1 and Pol I (Fig. 2j). Furthermore, ChIP-re-ChIP indicated that TBPL1 and Pol I are co-enriching especially at the IGS16 region (Fig. 2k), which harbors TCT motif on the DNA sense strand (Fig. 1i, Supplementary Fig. 4k). This prompted us to study the strand-specific effects of TBPL1 depletion on sincRNAs and asincRNAs at the IGS. We observed reductions in sincRNA and asincRNA levels at the IGS (Fig. 2l), suggesting that TBPL1 is required for the efficient function of Pol II and Pol I at the IGS. Consistent with this notion, TBPL1 depletion also lowered the enrichment of the Pol II-associated PAF1 across the IGS (Fig. 2m), indicating that TBPL1 is required for the efficient recruitment of Pol I, Pol II, and its associated factor PAF1.

Considering the interaction of TBPL1 with Pol I at the IGS, and the known regulation of ribosomal protein genes by TBPL1, we suspected that TBPL1 might be a master regulator of diverse transactional processes impacting ribosome biogenesis. So, we next asked whether TBPL1 also co-enriches with Pol I at the rRNA gene promoter. Indeed, ChIP-re-ChIP analysis revealed the co-enrichment of TBPL1 and Pol I at the rRNA gene promoter (Supplementary Fig. 4i). Furthermore, TBPL1-depleted cells showed a partial yet significant reduction in Pol I occupancy at the rRNA gene promoter (Supplementary Fig. 4j). Overall, our findings suggest that TBPL1promotes the localization and function of Pol I and Pol II at numerous genomic loci promoting ribosome biogenesis. Specifically, TBPL1 promotes Pol I loading at the rRNA gene promoter, Pol I and Pol II enrichment at the IGS, and Pol II recruitment to ribosomal protein genes. So, we propose that TBPL1 is a master regulator of ribosome manufacture.

### IGS Pol II transcription elongation via PAF1

Next, we aimed to further study the impact of PAF1 on the IGS. First, we set out to study the effect of PAF1 depletion on IGS ncRNA expression in the presence or absence of iPol I or iPol II. We used cells following a 48 h siRNA-mediated knockdown of PAF1 (Fig. 3a, b). PAF1 knockdown alone did not alter ncRNA levels within the IGS 16-18 region, from which IGS Pol I initiates ncRNA synthesis (Fig. 3c). However, PAF1 knockdown partly mimicked iPol I, decreasing IGS ncRNA levels in the IGS 20-30 region (Fig. 3c, Supplementary Fig. 5a). Pol I inhibition was dominant over PAF1 depletion, almost entirely repressing ncRNA levels across the IGS except for IGS16, where combining PAF1 knockdown with iPol I further decreased ncRNA levels (Fig. 3c). On another front, iPol II induced IGS ncRNA levels as expected (Fig. 3d). Notably, PAF1 knockdown failed to repress IGS ncRNA levels in the presence of iPol II despite the presence of readily detectable levels of ncRNAs (Fig. 3d). In fact, knocking down PAF1 slightly yet significantly increased IGS ncRNA levels at the IGS 16 and 30 sites. Together, these findings indicate that PAF1 maintains ncRNA levels in unperturbed cells. Also, PAF1 knockdown can moderately boost the ncRNA-repressing effect of iPol I and the ncRNA-inducing effect of iPol II. Strand-specific analysis of ncRNAs at the IGS further revealed that PAF1 knockdown decreases sincRNA and especially asincRNA levels (Fig. 3e). These results suggest that PAF1 promotes the function of Pol II and contributes to Pol I function at the IGS.

**Figure 3.**
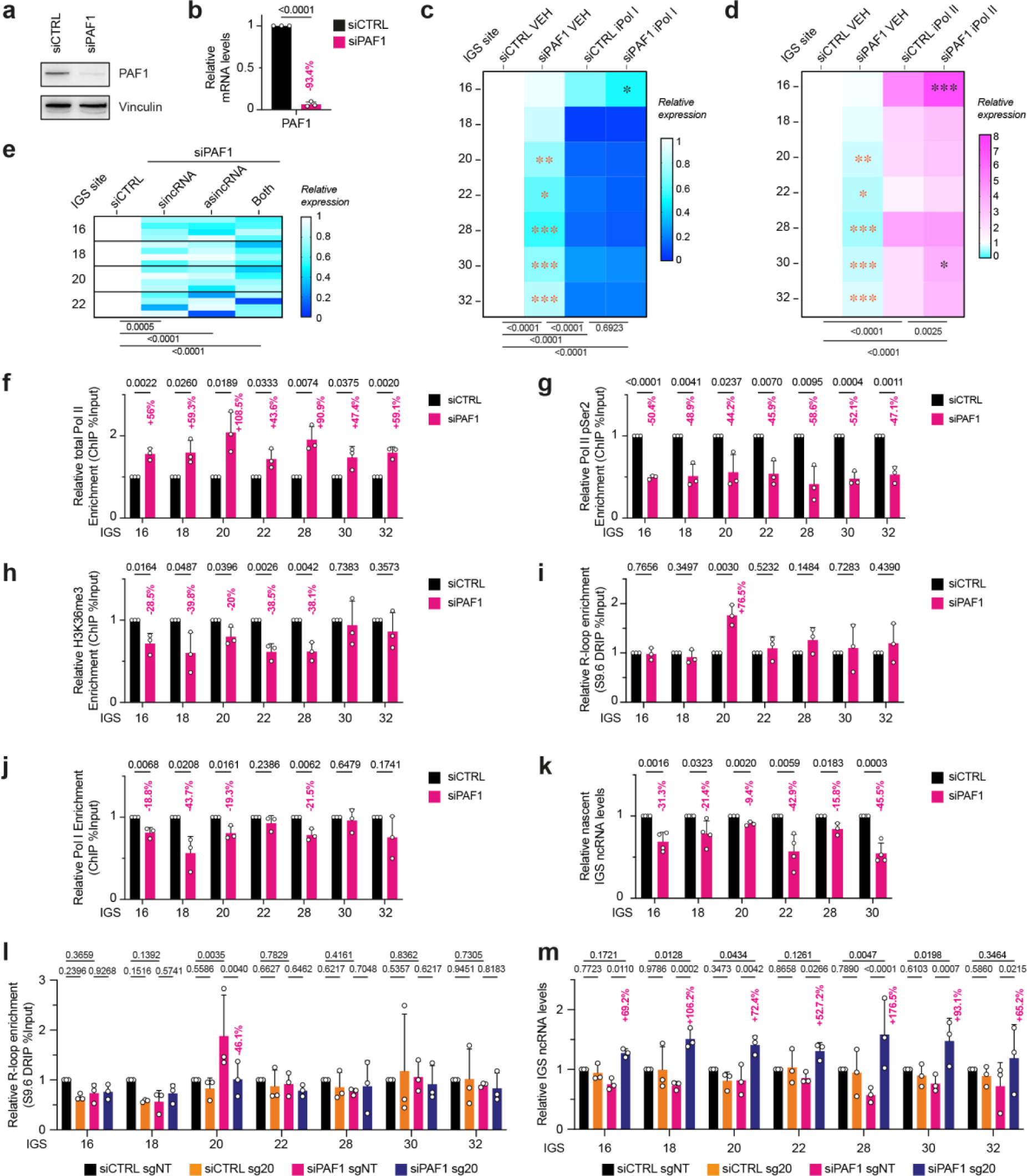
IGS-associated PAF1 mediates Pol II transcription elongation. (**a, b**) Validation of PAF1 knockdown via immunoblotting (**a**) and RT-qPCR (**b**). (**c, d**) Heatmaps showing the effect of PAF1 knockdown on IGS ncRNA levels in cells treated with vehicle (VEH), iPol I, or iPol II. (**e**) Heatmap presenting the strand-specific analysis of IGS ncRNA using ssRT-qPCR following PAF1 knockdown. **(f-h)** ChIP analysis assessing the impact of PAF1 knockdown on the enrichment of total Pol II (f), Pol II pSer2 (g), and H3K36me3 (h) across the IGS. **(i-k)** Impact of PAF1 knockdown on the enrichment of R-loops in DRIP (i), Pol I in ChIP (j), and nascent ncRNA levels in RT-qPCR (k) at the IGS. **(l, m)** Use of the LasR system employing the RED fusion protein together with sg20 represses R-loops at the IGS20 site in DRIP (l) and increases IGS ncRNA levels across the IGS in RT-qPCR (m). **(a-k)** Experiments were performed using HEK293T cells except for (k, m) where HEK293 TRex^TM^ were used; blots (a) are representative of three biologically independent experiments; quantitative data are shown as means (heat maps) or mean±s.d. (graphs); *n* = 3 except for (e) where *n* = 4 biologically independent experiments; data were analyzed using unpaired *t*-tests (b, f-k), two-way ANOVA with Dunnett’s multiple comparisons test (e), two-way ANOVA with uncorrected Fisher’s LSD test (l, m), and for (c, d) shown is two-way ANOVA with Tukey’s multiple comparisons tests (below heatmaps), multiple unpaired *t*-tests comparing VEH-treated samples (orange asterisks), and two-way ANOVA comparing iPol I- or iPol II-treated samples (black asterisks).

PAF1 can regulate transcription elongation within protein-coding genes by directly associating with Pol II and regulating histone marks, such as histone H3 lysine 36 trimethylation (H3K36me3)^26^. PAF1 knockdown led to the accumulation of total Pol II and depletion of Pol II pSer2, which represents elongating Pol II (Fig. 3f, g). These changes were observed in the absence of any change in Pol II protein level (Supplementary Fig. 5b). Decreases in H3K36me3 accompanied the changes in Pol II occupancy (Fig. 3h). These findings suggest that PAF1 regulates Pol II transcription elongation at the IGS.

Pol II stalling is linked to the formation of unscheduled R-loops^27,28,29^. Antisense R-loops created by an RNA polymerase can strongly obstruct RNA polymerases operating in the sense orientation at sites of convergent transcription^11^. Consistent with this notion, DRIP assays using the S9.6 antibody revealed the accumulation of R-loops specifically at IGS20 (Fig. 3i, Supplementary Fig. 5c), which was the site with the largest Pol II accumulation following PAF1 knockdown (Fig. 3f). S9.6 signals were reduced with *in vitro* RNaseH1 treatment, confirming S9.6 signal specificity (Supplementary Fig. 5c). This increase was further confirmed using GFP-dRNH1 (Supplementary Fig. 5d). The increase in R-loops at the IGS20 site was accompanied by losses in Pol I occupancy especially at the IGS 16-20 region (Fig. 3j) despite stable Pol I protein levels (Supplementary Fig. 5e). Consistent with the disruption of Pol I and Pol II at the IGS of PAF1-depleted cells, the latter displayed decreased ncRNA synthesis throughout the IGS (Fig. 3k). These findings suggest that PAF1 promotes IGS Pol II transcriptional elongation, preventing unscheduled R-loop accumulation at the IGS20 site. The accumulation of such unscheduled R-loops in PAF1-deficient cells may also limit Pol I function at the IGS.

To repress the unscheduled R-loops at IGS20, we employed the LasR system^11,12^. We used short guide RNAs (sgRNAs) targeting IGS20 (sg20) to selectively enrich the inducibly expressed RED (RNaseH1-EGFP-dCas9) fusion protein at the affected IGS site^11,12,30,31^. As controls, we used non-targeting sgRNA controls (sgNT) and the fusion protein dRED (dRNaseH1-EGFP-dCas9), which harbors a catalytically dead version of the RNaseH1 enzyme^11,12^. ChIP indicated the successful targeting of RED or dRED to IGS20 when using sg20 compared to sgNT (Supplementary Fig. 5f, g). DRIP analysis indicated the repression of the unscheduled R-loops at IGS20 in PAF1-depleted cells when targeting RED using sg20 (Fig. 3l). Resolving the unscheduled R-loops increased ncRNA levels throughout the IGS (Fig. 3m). Unlike RED, dRED did not lower R-loop levels or induce IGS ncRNA levels even when combined with sg20 (Supplementary Fig. 5h, i). Our findings suggest that the Pol II elongation defect-related accumulation of unscheduled R-loops at IGS20 in PAF1-depleted cells hinders baseline Pol I function at the IGS.

### Optimization of nucleoli via PAF1, TBPL1, and baseline sincRNAs

Inhibition of IGS Pol II or its dependent asincRNA-containing R-loops excessively induces Pol I-dependent sincRNAs^11^. The accumulating sincRNAs drastically disrupt nucleolar architecture and rRNA biogenesis^11^. Thus, we already know that Pol II-dependent asincRNAs promote ribosome biogenesis. In contrast, sincRNA accumulations are deleterious, and whether baseline sincRNA levels serve a biological function is unknown. Our findings suggest that TBPL1 and PAF1 operate through different mechanisms to promote the recruitment and function of Pol I and Pol II at the IGS. Thus, we hypothesized that baseline transcription by Pol I and Pol II maintains nucleolar organization and its role in rRNA biogenesis. Pre-rRNA synthesis occurs in the nucleolar FCs and at the FCs-DFC boundary before migrating through the DFC and GC as the transcript undergoes extensive sequential processing (Fig. 4a)^6,32^. Immunofluorescence and imaging analyses of the DFC-marking protein FBL and GC-indicating protein nucleophosmin (NPM) showed that the knockdown of PAF1 or TBPL1 partly changed the relative positioning of the DFC to the GC (Fig. 4b). Particularly, PAF1 or TBPL1 deficiency increased the levels of FBL-marked DFC bodies that protruded into the NPM-marked GC or were even located beyond it into the nucleoplasm. These phenotypes suggest that PAF1 and TBPL1 maintain the proper layering of nucleolar sub-compartments housing pre-rRNA synthesis and processing. This phenotype is distinct from the complete nucleolar fragmentation previously observed following the inhibition of Pol II or its R-loops using different approaches^11^.

**Figure 4.**
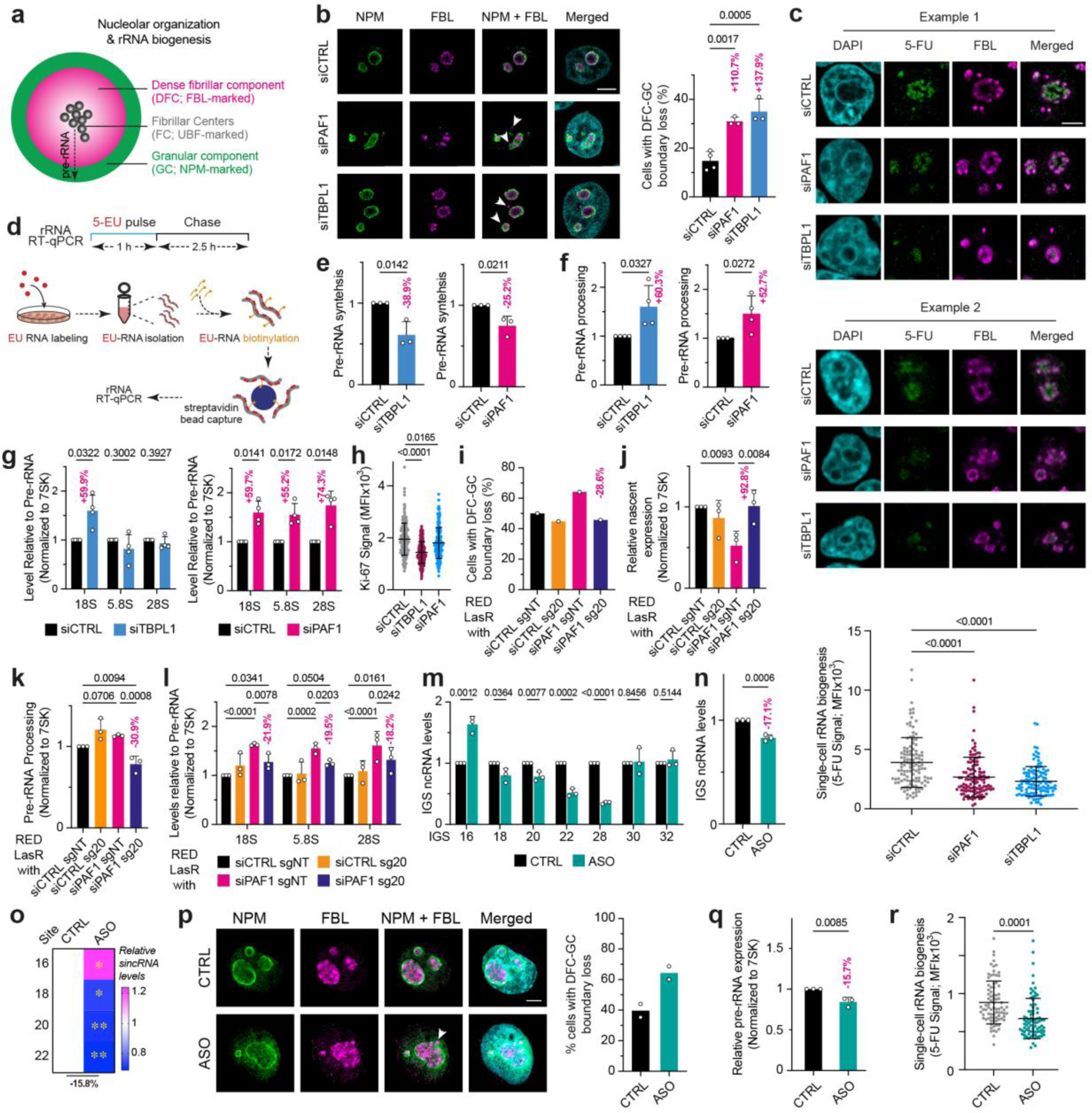
Nucleolar organization and function via PAF1, TBPL1, and sincRNAs. (**a**) Schematic of nucleolar organization and related rRNA biogenesis. (**b**) Knockdown of PAF1 or TBPL1 disrupts the boundary between the FBL-marked DFC and the NPM-marked GC. (**c**) Representative images and quantifications from single-cell analyses showing the impact of PAF1 or TBPL1 knockdown on rRNA biogenesis as assessed using 5-FU-labelled rRNA at the FBL-marked nucleolus. **(d-f)** Schematic (d) of the 5-EU-labeling and biotinylation-based pulse-chase RT-qPCR assay used to measure rRNA synthesis (e) and processing (f). (**g**) RT-qPCR analysis of the processing of the indicated mature rRNA molecules following the knockdown of PAF1 or TBPL1. **(h)** Impact of TBPL1 or PAF1 knockdown on Ki-67 signal. **(i-l)** Combining RED together with sg20 restores the DFC-GC boundary (h), restores rRNA synthesis (i) and decreases rRNA processing (j, k). **(m, n)** Impact of antisense oligonucleotides (ASOs) targeting sincRNAs between the IGS18-28 region on IGS ncRNA levels at single IGS sites (m) or all IGS sites (n) tested. **(o)** Impact of ASOs on sincRNA levels as assessed by ssRT-qPCR. **(p-r)** Impact of the ASOs on the nucleolar DFC-GC boundary (p), EU-labeled pre-RNA synthesis in RT-qPCR (q), and single-cell rRNA biogenesis assessed by 5-FU-labelled nucleolar rRNAs (r). **(b-r)** Experiments were performed using HEK293T cells except for (i-l) where 293 T-REx^TM^ were used; data shown as mean±s.d.; *n =* 3 (b-e, k-o, q, r), *n* = 2 (h), and *n* = 4 (f, g) biologically independent replicates; statistical analysis conducted using one-way ANOVA with Dunnett’s multiple comparisons test (b, h); one-way ANOVA with Sidak’s multiple comparisons test (c), unpaired *t*-tests (e-g, m-o, q, r), one-way (j) or two-way (l) ANOVA with Fisher’s LSD; one-way ANOVA with Tukey’s multiple comparisons tests (k); Images are representative of three (b, c) or two (p) independent experiments; Scale bar, 5 μm.

Therefore, we next set out to assess the potential role of PAF1 and TBPL1 in regulating rRNA biogenesis using 5-fluorouracil (5-FU), the incorporation of which into newly synthesized rRNAs can be quantified using single-cell microscopy. Notably, rRNA biogenesis was significantly disrupted in cells with TBPL1 or PAF1 knockdown (Fig. 4c). To assess the impact of TBPL1 and PAF1 on pre-rRNA synthesis and processing specifically, we next used 5-EU-labelled RNA pulse-chase assays coupled to RT-qPCR (Fig. 4d). Indeed, PAF1 or TBPL1 knockdown decreased pre-rRNA synthesis (Fig. 4e), consistent with the single-cell rRNA biogenesis results (Fig. 4c). We then assessed rRNA processing by quantifying the levels of 5-EU-marked 18S, 5.8S, or 28S rRNA relative to pre-rRNA levels following a 2.5-hour chase with unlabeled media (Fig. 4d). Notably, despite decreasing rRNA synthesis and rRNA biogenesis overall, the knockdown of PAF1 or TBPL1 increased pre-rRNA processing as assessed by 5-EU pulse-chase coupled to RT-qPCR (Fig. 4f). Of note, detailed analysis of the processing phenotype in PAF1-deficient cells indicated that processing related to all mature rRNAs was increased (Fig. 4g, right). This observation suggests an evenly accelerated processing rate of the different rRNA processing intermediates. In contrast, in TBPL1-deficient cells, the increased rate of rRNA processing was specific to 18S rRNA and was therefore not observed for the 28S or 5.8S rRNAs (Fig. 4g, left). These results suggest an asymmetrically increased rRNA processing in TBPL1-deficient cells. Consistent with the impact of TBPL1 or PAF1 on rRNA biogenesis, their depletion decreased cell growth as assessed by Ki-67 staining (Fig. 4h). Overall, as expected from our prior molecular studies (Fig. 2 and 3), these findings indicate that TBPL1 and PAF1 promote rRNA biogenesis through non-identical processes.

To examine whether the altered nucleolar organization and rRNA biogenesis in PAF1-depleted cells is dependent on the unscheduled IGS20 R-loops observed in these cells (Fig. 3i, l, m), we enriched the RED or control dRED fusion protein at IGS20 (Supplementary Fig. 5f, g). In PAF1-deficient cells, resolving the unscheduled R-loops at the IGS20 site (Fig. 3l), which moderately increases IGS ncRNA levels (Fig. 3m), restored nucleolar organization, rescuing rRNA synthesis, and re-restraining processing (Fig. 4 i-l). These findings suggest that low or baseline levels of ncRNAs may be critical for optimizing nucleolar organization and function.

Considering that Pol I-dependent sincRNAs constitute the bulk of baseline IGS ncRNAs^11^, we asked whether the direct repression of sincRNAs alters nucleolar organization and rRNA biogenesis, similar to the depletion of PAF1 or TBPL1. Therefore, we depleted the baseline levels of the Pol I-dependent sincRNAs in otherwise unperturbed cells using a pool of antisense oligonucleotides (ASOs) targeting sincRNAs synthesized from the IGS18-28 region (Fig. 4m). Notably, targeting sincRNAs in this region increased the levels of ncRNAs at IGS16, likely as part of a compensatory selection for cells with higher sincRNA transcription by Pol I initiating from IGS16 (Fig. 4m). Nonetheless, the ASO approach decreased the levels of IGS ncRNAs tested by an average of 17.1% (Fig. 4m). Indeed, strand-specific analysis confirmed that the sincRNA-targeting ASOs decreased relative sincRNA levels by an average of 15.8% across all sites tested (Fig. 4o). Notably, ASO-mediated knockdown of baseline sincRNAs partly phenocopied the loss of PAF1 or TBPL1, disrupting the DFC-GC nucleolar boundary (Fig. 4p). This defect in nucleolar organization was matched by a decreased rRNA biogenesis (Fig 4q, r). These findings suggest that a low level of Pol I-dependent sincRNAs is critical for nucleolar structure and function.

## Discussion

The nucleolus harbors the rDNA repeats encoding the rRNAs critical to universal protein synthesis^4,33^. At rDNA repeats, in addition to Pol I-dependent synthesis of pre-rRNA from rRNA genes, Pol I and Pol II transcribe ncRNAs from across the rDNA IGS. Previous work revealed a role for the IGS Pol II-dependent asincRNAs and R-loops in preserving nucleolar structure and function. Processes controlling Pol II function at the IGS and whether baseline IGS Pol I function serves a biological function was unclear. Our findings here reveal the regulation of Pol I and Pol II function at the IGS by TBPL1 and PAF1 through different processes and show that baseline Pol I function at the IGS optimizes rRNA biogenesis. Specifically, TBPL1 promotes the enrichment of Pol I and Pol II and consequent synthesis of sincRNAs and asincRNAs at the IGS (Fig. 5a). PAF1 also supports IGS Pol II elongation, preventing the formation of unscheduled R-loops that can hyper-repress Pol I enrichment and sincRNA synthesis (Fig. 5a). TBPL1 deficiency limits the enrichment of Pol I and Pol II, ultimately hyper-repressing sincRNA levels and compromising rRNA biogenesis (Fig. 5b). PAF1 deficiency leads to the accumulation of unscheduled R-loops that hyper-repress sincRNA levels and disrupt rRNA production (Fig. 5c). Together, our current and prior work^11^ indicate that the excessive depletion or accumulation of sincRNA levels disrupts nucleolar structure and function (Fig. 5d).

**Figure 5.**
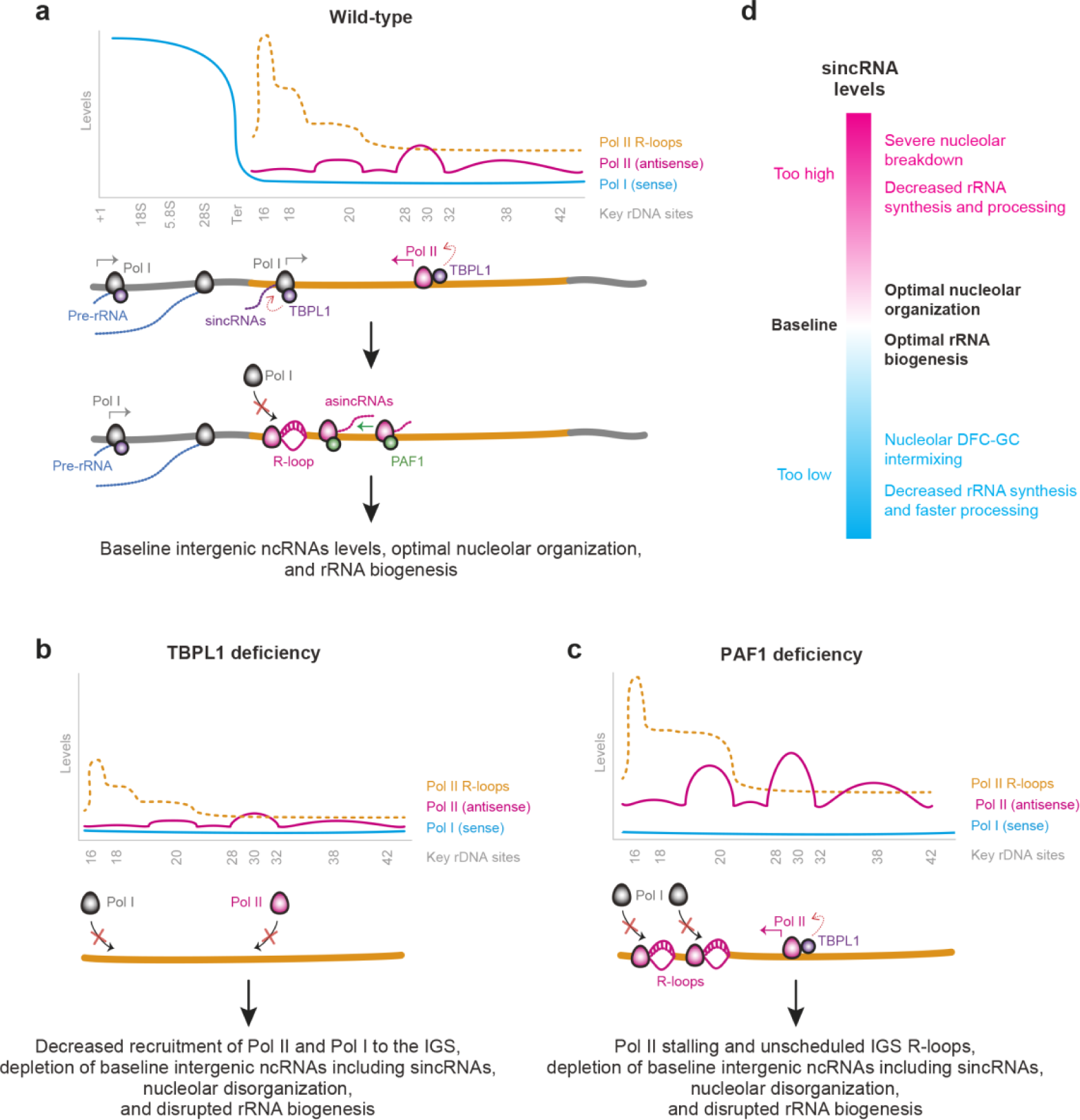
Model for the roles of Pol II, Pol I, and their ncRNAs at the rDNA IGS. **(a)** At the IGS of wild-type cells grown under standard conditions, TBPL1 at TCT motif-containing IGS sites promotes sense Pol I and antisense Pol II recruitment for the synthesis of sincRNAs and asincRNAs, respectively. IGS Pol II-associated PAF1 promotes the release of Pol II from a natural pause site at IGS20. Thus, TBPL1 and PAF1 promote the synthesis of baseline sincRNA levels through different processes. **(b)** TBPL1 deficiency reduces the recruitment and function of Pol I and Pol II at the IGS, decreasing the levels of intergenic ncRNAs and abrogating nucleolar structure and function. **(c)** Loss of PAF1 induces excessive Pol II pausing at IGS20, leading to unscheduled R-loops that further limit the ability of Pol I to synthesize the baseline levels of sincRNAs needed to ensure nucleolar organization and rRNA biogenesis. **(d)** Impact of sincRNA levels on nucleolar structure and function.

Pol II transcription initiation is regulated at multiple levels, including core promoter DNA sequences recognized by different transcription factors. TATA box is a common core promoter recognized by a TATA box-binding protein (TBP) that recruits other transcription factors, including TFIID and TAFs^34^. The *TBP* gene was duplicated with modifications during evolution, giving rise to different isoforms such as TBPL1, also known as TBP-related factor 2 (TRF2)^35^. TBPL1 preferentially recognizes TATA-less promoters with polypyrimidine motifs known as TCT promoters mostly at ribosomal protein genes^17,25^. Here, we found that TBPL1 is particularly enriched with the rDNA IGS at sites harboring such TCT motifs. Of note, TCT motifs are in the sense orientation at IGS sites where Pol I initiates sincRNA synthesis and in the antisense orientation at IGS sites from which Pol II initiates asincRNA synthesis. So, the positioning and orientation of these TCT motifs is consistent with our data indicating that TBPL1 promotes Pol I and Pol II recruitment and function at the IGS. Our results also revealed that TBPL1 promotes Pol I binding at the rRNA gene promoter. Together, our findings point to TBPL1 as a master regulator of coding or non-coding transcriptional units impacting rRNA manufacture.

Initiated Pol II commonly undergoes promoter-proximal pausing. Release of Pol II from pausing depends on the balance between DRB sensitivity inducing factors (DSIF), negative elongation factor (NELF), and positive elongation factor b (pTEFb), where DSIF and NELF stabilize Pol II in the paused state while pTEFb releases Pol II and promotes transcription elongation^36,37^. pTEFb, consisting of CDK9, is recruited to Pol II, where it phosphorylates DSIF, NELF and Ser2 on the CTD of Pol II itself^38,39^. PAF1 regulates this stage of Pol II transcription through the recruitment of chromatin remodeling and elongation factors. Here, we observed increased Pol II pausing and depletion of H3K36me3 in PAF1-deficient cells, suggesting that PAF1 promotes Pol II transcription elongation at the IGS. This is consistent with studies showing that PAF1C promotes Pol II elongation and the deposition of H3K36me3, H3K4me3, and H2Bub^26,40,41,42^. Other studies showed that PAF1C can promote Pol II pausing, but such contrasting findings may be linked to the different systems used or specific genes studied^43^. Nonetheless, in the absence of PAF1, we observed the accumulation of unscheduled R-loops at IGS20, which can result from excessive or longer Pol II pausing^27,28,29,40^. Notably, the processes uncovered in our study here may also impact tumorigenesis. For instance, cells lacking the BRCA1 tumor suppressor rely on the ability of RNF8 and RNF168 to ubiquitylate and recruit the R-loop-repressive factors XRN2 and DHX9, respectively, to prevent the excessive accumulation of R-loops at the rDNA IGS^44,45^. Excessive R-loop accumulation in BRCA1-mutant cells deficient in RNF8 or RNF168 stalls cellular growth and prevents tumor growth^44,45^.

In response to environmental stresses such as heat shock, cells repress asincRNAs and induce the expression of certain sincRNAs^11,46,47,48,49^. Under such stress conditions, the induced sincRNAs may promote the transition of the nucleolus from a liquid-like to solid-like structure as part of a process that promotes cell survival under stress^11,46,47,50,51^. Our data here show that baseline sincRNA levels can exert beneficial effects, even in the absence of environmental stress. Notably, under heat shock, the abundance of various proteins controlling Pol II function increases in the nucleolus^52^. We expect future studies to assess the impact of PAF1, TBPL1, and other Pol II regulatory factors on the expression and function of sincRNAs under various conditions including during the activation of the nucleolar stress response as in during heat shock. Also, under heat shock conditions, a long ncRNA called PAPAS can be transcribed in the antisense orientation to the rRNA gene promoter^50^. PAPAS may promote the heterochromatinization of the rRNA gene promoter under stress^50^. Also, the ‘ncRNA’ PAPAS can encode for a protein called Ribosomal IGS-encoded protein (RIEP)^53^. Of note, RIEP enriches mostly at the Pol II IGS promoter region^53^. Thus, it will be critical to assess whether the heat-shock-dependent enrichment of RIEP intersects with TBPL1 and PAF1 function to modulate IGS Pol II activity and Pol I-dependent sincRNA synthesis under stress.

Inhibition of Pol II using flavopiridol (inhibits CDK9 phosphorylation of Pol II CTD) or α-amanitin (degrades Pol II) hyper-induces sincRNAs, robustly disrupting nucleolar organization and ribosome biogenesis^11^. In the presence of Pol II inhibition, partly repressing the hyper-induced sincRNAs using ASOs partially rescues nucleolar organization and rRNA production^11^. Such partial rescue could be due to the incomplete repression of sincRNAs. Partial rescue only in the context of CDK inhibition may also be tentatively linked to the potential presence of other nucleolar CDK targets^54^.

Like all studies, our work here has some limitations. First, we have identified a number of PICs co-enriched with nucleolar Pol II. MED26 and DR1 were validated to interact with Pol II and to be nucleolar, but we could not detect their enrichment at rDNA. Such PICs may interact with nucleolar Pol II off chromatin or might co-enrich with Pol II onto chromatin at rDNA under specific conditions. We also note that the broader list of hits from the compBioID may also include some factors that interact with Pol II in the nucleoplasm before localizing to the nucleolus. So, future work focusing on nucleolar Pol II-linked PICs from this study should similarly validate their interaction with Pol II, nucleolar localization, and co-enrichment with Pol II at rDNA or even perinucleolar chromatin. Second, in addition to the targeted repression of IGS R-loops, the exogenous expression of sincRNAs would have been helpful as an additional tool to rescue defects in TBPL1 or PAF1 deficient cells. However, rescuing phenotypes via such exogenous sincRNA overexpression presents several challenges. These include determining the exact levels and mixtures of distinct sincRNAs needed and achieving perfect nucleolar and sub-nucleolar localization, amongst other problems. We rescued nucleolar structure and function by selectively ablating the unscheduled IGS R-loops in PAF1-deficient cells. So, even if PAF1 can impact myriad processes throughout the nucleus, the targeted ablation of the unscheduled R-loops in PAF1-deficient cells allowed us to isolate the critical molecular event required for the disruption of nucleolar structure and function in PAF1-deficient cells. Third, due to the ability of TBPL1 to impact numerous processes linked to ribosome biogenesis, we could not devise a rescue experiment to rescue rRNA biogenesis in TBPL1-deficient cells. Fourth, other limitations of our study include those linked to using the RED/dRED-based LasR system for targeted R-loop modulation, which we have described in detail previously^12^. For instance, all limitations associated with CRISPR- and dCas9-based technologies apply to this system, including poorly understood yet rare chromosome rearrangements^55^. Fifth, rDNA repeats are highly repetitive and so our analysis of chromatin at such loci represents an average picture of the state of the repeats across the cell population and should not be interpreted as absolute measurements related to any single rDNA unit, rRNA gene or IGS region.

Overall, we report the first interactome of nucleolar Pol II. This interactome helped reveal the regulation of Pol II and Pol I by TBPL1 and PAF1 proteins at the rDNA IGS and the beneficial impact of these two proteins on nucleolar structure and function. We also uncovered the role of Pol I and baseline sincRNAs at the IGS in maintaining rRNA manufacture. Our findings will have important implications for our understanding of cell growth in the presence and absence of environmental stresses and in the context of physiological and pathological settings such as cancer. Finally, we anticipate that the nucleolar Pol II interactome uncovered here will continue yielding critical insights into rRNA biogenesis and other processes in the future.

## Methods

### Cell culture

Human embryonic kidney 293 cells (HEK293T) were maintained in Dulbecco’s modified Eagle’s medium (DMEM, Wisent Bioproducts) supplemented with 10% (*v/v*) of fetal bovine serum (FBS, Wisent) and 1% (*v/v*) penicillin and streptomycin (Wisent) at 37°C and 5% CO_2_ condition. 293 T-REx^TM^ cells (ThermoFisher Scientific) were maintained in DMEM supplemented with 10% (*v/v*) tetracycline-free FBS and 1% penicillin and streptomycin. Flp-In 293 T-REx cells grown in DMEM supplemented with 10% FBS (Wisent) were used to generate BioID cell lines to interrogate interacting partners of the RNA Pol II subunit POL2RI/RPB9, whose ORF was cloned into a pcDNA5 FRT/TO 3xFLAG MiniTurboID plasmid (Raught Lab)^56^ such that the biotin ligase fusion is N-terminal to the protein. Stable T-REx lines were generated by transfecting 1 μg of pcDNA5 FRT/TO 3xFLAG MiniTurboID and 2 μg of pOG44 (Invitrogen, Cat# V600520) with Lipofectamine3000 (Invitrogen, Cat # L3000001) in a 6-well plate at 60% confluency. Cells were passaged into a 10 cm dish and subsequently selected for resistance to 200 μg/mL hygromycin B (Gibco). For knockdown experiments, HEK293T cells at 50% confluence were transfected with 50 nM siRNA using Lipofectamine RNAiMAX (Invitrogen) as described by the manufacturer in an antibiotic-free growth medium for 48 h. For inhibitory experiments, 2 µM flavopiridol (Santa Cruz Cat # sc-202157; iPol II), 50 ng/ml actinomycin D (LAD/iPol I; Sigma Cat# A1410-2MG), or DMSO vehicle control were used. For locus-associated targeting of RED or dRED as part of the LasR system, HEK293T T-REx^TM^ cells were grown at 70% confluence and used as described previously^12^. For knockdown experiments, cells were transfected using Lipofectamine RNAiMAX (Invitrogen, Cat# 13778075). Antisense oligonucleotides (ASOs) used to knock down sincRNAs were as follows. We used the antisense LNA GapmeR negative control B (Cat# 339515, LG0000001-DDA; gctcccttcaatccaa), IGS18 (LG00210930-DDA; agtgtgctctgtgaac), IGS20 (LG00210936-DDA; acgcaagaaaggaaga), IGS22 (LG00210956-DDA; acgtgaccgagagaaa) and IGS24 (LG00210966-DDA; gtgacgtgtagagatt). Commercially available siRNAs were used for TBPL1 (Dharmacon, L1:L-017254-00-0005) and its control (Dharmacon, D-001810-10-05), and PAF1 (Thermo Fischer, s29267) and its control (Thermo Fischer, Cat# 4390843).

### BioID

Proximity-dependent biotinylation (BioID^57^) was performed as previously described^58^. Briefly, stable isogenic cell pools expressing FmT-alone (“tag only” controls) or the FmT-fused bait protein(s) were expanded to 5 x 15 cm^2^ plates at ∼80% confluency. Expression of the fusion protein was induced by the addition of 1 ug/ml tetracycline to the culture media for 24 h. The following day, cells were treated with 50 μM biotin (final concentration) for 60-90 min. Cells were collected by scraping, and pelleted (2,000 rpm, 3 min). The pellet was washed twice with PBS, and dried pellets snap frozen. Cell pellets were thawed on ice and resuspended in 10 ml lysis buffer (50 mM Tris-HCl pH 7.5, 150 mM NaCl, 1 mM EDTA, 1 mM EGTA, 1% Triton X-100, 0.1% SDS, 1:500 protease inhibitor cocktail (Sigma-Aldrich), 1:1,000 benzonase nuclease (Novagen)), then incubated on an end-over-end rotator at 4°C for 1 h, briefly sonicated to disrupt any visible aggregates, and centrifuged at 45,000 x g for 30 min at 4°C. The supernatant was transferred to a fresh 15 ml conical tube. 30 μl of packed, pre-equilibrated (in lysis buffer) streptavidin-sepharose beads (GE) were added and the mixture was incubated for 3 h at 4°C with end-over-end rotation. Bound beads were pelleted by centrifugation at 2,000 rpm for 2 min and transferred with 1 ml of lysis buffer to a fresh Eppendorf tube, washed once with 1 ml lysis buffer and twice with 1 ml of 50 mM ammonium bicarbonate (ammbic, pH 8.3), transferred in ammbic to a fresh centrifuge tube and washed two more times with 1 ml ammbic. Tryptic digestion was performed by incubating beads with 1 μg MS-grade TPCK trypsin (Promega, Madison, WI) dissolved in 200 μl of 50 mM ammbic (pH 8.3) overnight at 37°C. The following morning, an additional 0.5 μg MS-grade TPCK trypsin was added, and the beads were incubated an additional 2 h at 37°C. The beads were pelleted by centrifugation at 2,000 x g for 2 min, and the supernatant was transferred to a fresh Eppendorf tube. Beads were washed twice with 150 μl of 50 mM ammbic, and washes pooled with the first eluate, to collect liberated tryptic peptides. The peptide mix was lyophilized and resuspended in buffer A (0.1% formic acid). 1/6th of the sample was analyzed per MS run.

Validation of biotinylation was conducted using HRP Streptavidin (Abcam ab7403) western blots. For whole cell and nucleolar fraction BioID experiments, five and ten 15 cm^2^ cell culture dishes were used, respectively. Membranes were blocked for 1 h at room temperature with 3% BSA in TBST and subsequently incubated with 1:15,000 dilution of streptavidin HRP in blocking buffer for 30 min at room temperature.

### Mass spectrometry

High performance liquid chromatography was conducted using a 2 cm pre-column (Acclaim PepMap 50 mm x 100 um inner diameter (ID)), and 50 cm analytical column (Acclaim PepMap, 500 mm x 75 um ID; C18; 2 um; 100 Å, Thermo Fisher Scientific, Waltham, MA), running a 120 min reversed-phase buffer gradient at 225 nl/minute on a Proxeon EASY-nLC 1000 pump in-line with a Thermo Q-Exactive HF quadrupole-Orbitrap mass spectrometer. A parent ion scan was performed using a resolving power of 60,000, then up to the twenty most intense peaks were selected for MS/MS (minimum ion count of 1,000 for activation) using higher energy collision induced dissociation (HCD) fragmentation. Dynamic exclusion was activated such that MS/MS of the same *m/z* (within a range of 10 ppm; exclusion list size = 500) detected twice within 5 sec were excluded from analysis for 15 sec. For protein identification, Thermo .RAW files were converted to the .mzXML format using Proteowizard^59^, then searched using X!Tandem^60^ and COMET^61^ against the Human RefSeq Version 45 database (containing 36,113 entries). Data were analyzed using the trans-proteomic pipeline (TPP)^62^ via the ProHits software suite (v3.3)^63^. Search parameters specified a parent ion mass tolerance of 10 ppm, and an MS/MS fragment ion tolerance of 0.4 Da, with up to two missed cleavages allowed for trypsin. Variable modifications of +16@M and W, +32@M and W, +42@N-terminus, and +1@N and Q were allowed. Proteins identified with an iProphet cut-off of 0.9 (corresponding to ≤1% FDR) and at least two unique peptides were analyzed with SAINT Express v.3.6^64^. Twenty control runs (from cells expressing the FlagBirA* epitope tag alone) were collapsed to the four highest spectral counts for each prey and compared to the experimental data, consisting of two biological replicates (each analyzed with two technical replicates). High confidence interactors were defined as those with bayesian false discovery rate (BFDR) ≤0.01.

### Sucrose gradient-based assays

Cells grown to 70-80% confluence in 15 cm plates were washed twice with S1 solution before scraping the cells using S1 solution (1 mM Sucrose, 3 mM MgCl_2_) and spinning at 1,000 rpm for 10 min at 4°C. The cell pellet was resuspended with 500 μL of S1 solution and a small aliquot was saved as the whole cell fraction sample, before sonicating five times at 50% for 10 sec each. 700 μL of S2 solution (1 M Sucrose, 3 mM MgCl_2_) was then added to the bottom of the tube before spinning at 1.8 r.c.f. for 10 min at 4°C. A small aliquot of the supernatant was saved as the nucleoplasmic/cytoplasmic fraction sample before all the supernatant was discarded. The nucleolar pellet was then resuspended in S1 solution, and more rounds of fractionation were performed depending on the size of the pellet. Samples were then resuspended in lysis buffer (50mM Tris pH8, 120mM NaCl, 0.5% NP-40, 1X protease inhibitor) and needle-shredded before running on SDS-PAGE or analysis by mass spectrometry.

### RNA extraction

Total RNA was isolated from cells using a RNeasy mini kit (Qiagen, Cat# 74104) as described in the manufacturer’s protocol. Reverse transcription was performed as previously described^11^. In summary, 1 μg of total RNA was treated with DNase I followed by reverse transcription using dNTPs, random hexamers, 5X first-strand buffer, DTT, RNaseOUT, and MMLV reverse-transcriptase at 25°C for 10 min, 37°C for 60 min and 70°C for 15 min. The cDNA was diluted 1:10, and a 4 µL sample was used for qPCR-based analyses. qPCR reactions were performed at 95°C for 5 min and 60°C for 30 sec, followed by 39 cycles of 95°C for 5 sec and 60°C for 30 sec.

### Strand-specific RT-qPCR (ssRT-qPCR)

We used previously designed primers and followed as described previously. Briefly, separate 10μL reverse transcription reactions were set up, where 200ng of purified RNA was incubated with 2.5nM strand-specific tagged primer, 2.5nM control strand-specific primer (for example, 7SK), 1mM dNTPs, 1x first-strand buffer, 10mM DTT, 40 U RNaseOUT, and 200 U M-MLV reverse transcriptase. For each RNA sample, false-primer reactions were carried out using DEPC ddH_2_O instead of the transcript of interest. Reactions were carried out in the same cycling as above. cDNA samples were diluted 1:10 and amplified using strand-specific primers at the IGS and 7SK. qPCR reactions were performed as above.

### Population-level pulse-chase assays

Cells cultured to a 50% confluency in a 6-well plate were pulsed with 20 mM 5-EU for 1 h, followed by either RNA extraction or a 2.5 h chase using non-labeled growth media. Total RNA was isolated as described above in the ‘RNA extraction’ section. Click-iT Nascent RNA capture kit (Invitrogen, Cat# C10365) was used as described in the manufacturer’s protocol to isolate 5-EU-ladled RNA. In summary, 1 μg of total RNA was incubated with Click-iT EU buffer, CuSO_4_, biotin azide, Click-iT reaction buffer additive 1 for 3 min before the addition of Click-iT reaction buffer additive 2 and incubation for 30 min at room temperature. The RNA was precipitated by incubating overnight with UltraPure Glycogen, 7.5 M ammonium acetate, and 100% ethanol at -80°C, followed by centrifugation at 13,000 *g* for 20 min at 4°C and two rounds of washes with 75% (*v/v*) ethanol. 0.75 µg of the precipitated RNA was then incubated with 2X Click-iT RNA binding buffer and RNaseOUT for 5 min at 67-70°C followed by the addition of streptavidin beads and incubation for 30 min at room temperature with frequent agitation. The RNA beads were washed five rounds with Click-iT reaction wash buffer 1 followed by wash buffer 2. The beads were then incubated at 68-70°C for 5 min followed by reverse transcription and qPCR as described above.

### Single-cell rRNA biogenesis assays

Cells previously seeded and transfected in 24-well plates were used. On the day of the assay, the cells were exposed to 1 mM 5-fluorouracil (5-FU; Sigma, Cat# F5130) for 15 min, followed by a gentle wash with unlabelled media. Cells were then fixed, permeabilized, and stained as described below (see the ‘Immunofluorescence’ section). Staining was performed using antibodies for BrdU (1:500), FBL (1:500), or POLR1A (1:500) antibodies.

### Immunofluorescence

25,000-50,000 cells were seeded on poly-L-lysine coated coverslips for 24 h. Cells were then fixed using 4% (*v/v*) paraformaldehyde for 15 min, followed by three washes with 1X PBS. Cells were then permeabilized with 0.5% (*v/v*) Triton X-100 for 15 min, followed by three washes with 1X PBS before incubation with 5% (*w/v*) bovine serum albumin (BSA) for 1 h at room temperature. The cells were then incubated with the primary antibody in 5% BSA overnight at 4°C in a humidified chamber. The following day, coverslips were washed thrice with 1X PBS and incubated with the secondary antibody at 1:500 (*v/v*) in 5% BSA. Coverslips were then incubated with 1X DAPI for 5 min before mounting on slides using DAKO mounting media (Agilent Technologies, Cat #S302380-2). Images were captured at 60X or 100X using a Nikon C2+ confocal microscope coupled to NIS-Elements AR software (Nikon).

### Co-immunoprecipitation

Cells at a 70-80% confluency were washed twice with ice-cold 1X PBS before being lysed using 1X CoIP lysis buffer (50 mM Tris pH 7.5, 150 mM NaCl, 2.5 mM MgCl_2_, 1 mM DTT, 1% Triton X-100, 1X protease inhibitor). Lysates were incubated for 30 min with 5 μL of Benzonase under agitation at 4°C. Lysates were precleared with magnetic Dynabeads protein A (Invitrogen, Cat# 10006D) for 1 h at 4°C while 2.5 µg of the antibody was incubated with 10 µL of magnetic beads for 2 h at 4°C. Then 2.5 mg of the precleared lysate was incubated with the antibody-bead complex for 1 h at 4°C. The immunocomplex was washed thrice with 1X PBST and eluted off the beads using 50 µL of 2X SDS loading buffer at 95°C for 5 min. Samples were then analyzed by SDS-PAGE.

### Chromatin immunoprecipitation (ChIP)

Cells grown to 70-80% confluency were crosslinked with 1% formaldehyde for 10 min, followed by 5 min incubation with 2.5 mM glycine to quench the reaction. Cells were lysed using lysis buffer (5 mM PIPES, 85 mM KCL, 0.5% (*v/v*) NP-40, 1X protease inhibitor), incubated on ice for 10 min followed by a 10 min spin at 10,000 r.p.m. and 4°C. The cell pellet was incubated with nuclear lysis buffer (50 mM Tris-HCL, 10 mM EDTA, 1% (*w/v*) SDS, 1X protease inhibitor) for 10 min on ice followed by eight rounds of sonication for 20 sec each at 40% amplitude. Samples were then spun at 12,500 *g* for 10 min at 4°C. 50 µL of the sheared chromatin was used for the immunoprecipitation by dilution at 1:10 (*v/v*) in IP dilution buffer (16.7 mM Tris-HCL pH 8.0, 0.01% (*w/v*) SDS, 167 mM NaCl, 1.2 mM EDTA, 1.1% (*v/v*) Triton X-100, and 1X protease inhibitor) and incubation with 2-5 µg of the given antibody overnight at 4°C with constant rotation. 20% of the sheared chromatin was also aliquoted and stored at 4°C. The immunocomplex was then incubated with pre-washed magnetic Dynabeads protein A (Invitrogen, Cat# 10006D) for 2 h at 4°C with constant rotation. The bead-immunocomplexes were then washed with low salt buffer (20 mM Tris-HCl, 0.1% (*w/v*) SDS, 1% (*v/v*) Triton X-100, 2 mM EDTA, 150 mM NaCl), high salt buffer (20 mM Tris-HCl, 0.1% (*w/v*) SDS, 1% (*v/v*) Triton X-100, 2 mM EDTA, 500 mM NaCl), LiCl wash buffer (10 mM Tris-HCl, 1% (*v/v*) NP-40, 1% (*w/v*) sodium deoxycholate, 1 mM EDTA, 250 mM LiCl), and two rounds of TE buffer (10 mM Tris-HCl pH 8.0, 1 mM EDTA) before incubation twice with elution buffer (1% (*w/v*) SDS, 100 mM NaHCO_3_) for 15 min each. The eluates were de-crosslinked by incubating overnight at 65°C with 200 mM NaCl. The RNA and proteins were depleted from the samples using a 30-min incubation with RNase A and two 2-hour incubations with Tris-HCl, EDTA, and proteinase K at 45°C. The DNA was then purified using the Gel/PCR DNA-fragment extraction (Geneaid, Cat# DF3000) and diluted at 1:5 before qPCR-based analyses.

### Sequential ChIP (ChIP-re-ChIP)

Before beginning, antibodies to be used for the first round of immunoprecipitation were bound and crosslinked to Protein A/G magnetic beads using Pierce^TM^ Crosslink Magnetic IP/Co-IP Kit (Thermo Fischer, Cat# 88805) following the manufacturer’s instructions. Briefly, 25 µL of magnetic beads were washed and incubated with 10 µg of antibody for 15 min at room temperature. Samples were then crosslinked by incubating with 20 µM DSS for 30 min at room temperature. The antibody-bead mixture was then washed twice with elution buffer followed by IP buffer. ChIP-re-ChIP was performed similarly to ChIP (see above) but with some modifications. The sheared chromatin is incubated with 5µg of antibody crosslinked to beads overnight at 4°C. The immunocomplex was washed as done in regular ChIP before eluting off the beads using 50 µL of Elution buffer (pH 2.0) from Pierce^TM^ Crosslink Magnetic IP/Co-IP for 5 min. Samples were then neutralized using 4 µL of Neutralization Buffer (pH 8.5) from Pierce^TM^ Crosslink Magnetic IP/Co-IP Kit before dilution at 1:10 in IP dilution as in ChIP and incubating with 2-5 µg of antibody overnight at 4°C for the second round of immunoprecipitation. The remaining steps in the protocol were the same as described in the ‘ChIP’ section above.

### DNA-RNA hybrid immunoprecipitation (DRIP)

DRIP experiments were performed as previously described^11^. In summary, cells at 70-80% confluency in 6 cm plates were washed twice with 1X PBS. Cell pellets were resuspended in TE buffer and incubated overnight at 37°C with 0.5% SDS and proteinase K for cell lysis and protein digestion. Genomic DNA was isolated using phenol-chloroform followed by precipitation with 3M NaOAc pH 5.2 and 100% ethanol. The DNA was spooled out and washed with 70% ethanol before it was left to dry. The isolated DNA was then resuspended in TE buffer and incubated overnight at 37°C with HindIII (New England Biolabs (NEB), Cat# R01045), EcoRI (NEB, Cat# R0101L), BsrgI (NEB, Cat# R05755), XbaI (NEB, Cat# R01455), SSPI (NEB, Cat# R0132), buffer 2.1, and spermidine. The digested DNA was then purified using 3 M NaOAc pH 5.2, phenol-chloroform, and 100% ethanol before washing it with 70% ethanol and leaving it to dry. The precipitated DNA was resuspended with TE (10 mM Tris-HCl pH 8.0, 1 mM EDTA) buffer. Then, 4.4 µg of DNA was incubated with RNaseH1 overnight at 37°C before incubating with 10 µg of IgG or S9.6 antibody overnight at 4°C. The antibody-DNA complexes were incubated with protein G magnetic beads for 2 h at 4°C, then washed thrice with 1X binding buffer. Samples were then eluted off the beads by incubating with DRIP elution buffer and proteinase K for 45 min at 55°C. The DNA was then purified using Gel/PCR DNA-fragment extraction (Geneaid, Cat# DF3000) before qPCR-based analyses.

### Statistical analysis

Statistical tests are indicated in the figure legends. Most of the statistical analysis consisted of unpaired t-tests or one-way or two-way ANOVA with multiple comparisons tests including Dunnett’s, Sidak’s, and Tukey’s. Individual P values are indicated for all data. For heatmaps, statistical significance is indicated as *p<0.05, **p<0.01, ***p<0.001, or ****p<0.0001. For mass spectrometry, PICs with a false discovery rate (FDR) ≤1% were retained.

## Acknowledgments

We thank members of the Mekhail and Hakem labs for fruitful discussions and feedback. We also thank Dr. Karlene Cimprich for the RNaseH1(DN) tool. N. Khosraviani was supported by a Vanier Doctoral Scholarship from the Canadian Institutes of Health Research (CIHR). R. Krishnan was supported by the Princess Margaret Cancer Foundation and the Hold’Em for Life Foundation. Work that is related to this study was supported by CIHR funds (PJT 183597 and 18607) to R. Hakem. This study was mainly supported by CIHR funds (PJT 159676 and 190143) to K. Mekhail, who also acknowledges support from the Royal Society of Canada.

## Author contributions

N.K. and K.M. wrote the text, which all co-authors edited. N.K. and K.M. designed most experiments except for the compBioID experiments, which were designed by V.T.Y., J.St-G., B.R., and K.M. All experiments were conducted by N.K. with assistance from Y.Y.H. to ChIP, RNA, and image analysis, from S.B.C. and C.G. to RNA analysis, and from R.K. to the RNH1-based R-loop measurements, except for the compBioID experiments, which V.T.Y. and J.St-G performed. R.H. supervised R.K. and B.R. supervised J.St-G. while N.K., V.T.Y., J.St-G., Y.Y.H., S.B.C., and C.G. were supervised by K.M.

