## Supplementary Information for "Nucleolar Pol II interactome reveals TBPL1, PAF1, and Pol I at intergenic rDNA drive rRNA biogenesis"

### Supplementary Information (SI) for Khosraviani *et al.*

**SI content summary:**

Supplementary tables

Supplementary figure legends

Supplementary figures

Supplementary Dataset (attached .xlsx file)

**Supplementary Table 1. List of antibodies used in this study.**

| Antibody | Company | Catalogue Nb. | Experiment |
|---|---|---|---|
| IgG (Rb) | Abcam | ab171870 | ChIP (2.5 μg), IP (2.5 μg) |
| IgG (Ms) | Millipore | 12-371 | ChIP (2.5 μg), IP (2.5 μg) |
| Pol II CTD (Ms) | Cell Signaling | 2629S | ChIP (1:100), IP (1:100) |
| Pol II pSer2 (Rb) | Abcam | ab5095 | ChIP (5 μg) |
| PolR1A (Rb) | Cell Signaling | 24799S | ChIP (1:100), IP (1:100) |
| Fibrillarin (Rb) | Abcam | ab5821 | IF (1:250) |
| B23 (Ms) | Sigma | B0556 | IF (1:500) |
| PAF1 (Rb) | Abcam | ab20662 | WB (1:1000) ChIP (5 μg), IP (2.5 μg) |
| MED26 (Rb) | Cell Signaling | 149505 | WB (1:1000), ChIP (1:100), IP (1:100) |
| TBPL1 (Rb) | Proteintech | 12258-1-AP | WB (1:1000), ChIP (5 μg), IP (2.5 μg) |
| H3K36me3 | Abcam | ab9050 | ChIP (2.5 μg) |
| Vinculin (Ms) | Sigma | V9131 | WB (1:3000) |
| DR1 (Rb) | Cell Signaling | 6447S | WB (1:1000), ChIP (1:100) |
| BrdU (Ms) | Sigma | B2531 | IF (1:100) |
| LEO1 | Proteintech | 12281-1-AP | WB (1:1000) |
| CDC73 | Cell Signaling | 8126S | WB (1:1000) |
| LDHA | Sigma | MABC150 | WB (1:1000) |
| S9.6 | | In-house (S9.6 Hybridoma cells) | DRIP (10μ) |
| HRP Streptavidin | Abcam | ab7403 | WB (1:15,000) |
| GFP | Abcam | ab290 | ChIP (5 μL) |
| Ki67 | eBioscience | SolA15 | IF (1:500) |

**Supplementary Table 2. List of primer sequences used in this study.**

| Primer | Sequence | Application |
|---|---|---|
| IGS 16 F | ACACACACACACCCCGTAGT | ChIP-qPCR, RT-qPCR, DRIP-qPCR |
| IGS 16 R | GAAATGGGGCTTCGATACAT | ChIP-qPCR, RT-qPCR, DRIP-qPCR |
| IGS 18 F | GTTGACGTACAGGGTGGACTG | ChIP-qPCR, RT-qPCR, DRIP-qPCR |
| IGS 18 R | GGAAGTTGTCTTCACGCCTGA | ChIP-qPCR, RT-qPCR, DRIP-qPCR |
| IGS 20 F | GTAGCCTTGGGCTTCTCTCC | ChIP-qPCR, RT-qPCR, DRIP-qPCR |
| IGS 20 R | AGTTTTCAGCCCCAACACAC | ChIP-qPCR, RT-qPCR, DRIP-qPCR |
| IGS 22 F | CAGTGGCTCACGTCTGTCAT | ChIP-qPCR, RT-qPCR, DRIP-qPCR |
| IGS 22 R | CGCCTGACTCCATTTCGTAT | ChIP-qPCR, RT-qPCR, DRIP-qPCR |
| IGS 28 F | CCTTCCACGAGAGTGAGAAG | ChIP-qPCR, RT-qPCR, DRIP-qPCR |
| IGS 28 R | GACCTCCCGAAATCGTACAC | ChIP-qPCR, RT-qPCR, DRIP-qPCR |
| IGS 30 F | GGTCTCTGCGTCTCGCTATC | ChIP-qPCR, RT-qPCR, DRIP-qPCR |
| IGS 30 R | TGAAGAATTCAGGCCTTGGT | ChIP-qPCR, RT-qPCR, DRIP-qPCR |
| IGS 32 F | AAAAGCTGGCCGATCTGAAT | ChIP-qPCR, RT-qPCR, DRIP-qPCR |
| IGS 32 R | CGTCTGTTCAGCTATTTTGCAG | ChIP-qPCR, RT-qPCR, DRIP-qPCR |
| Pre-rRNA F | GGCGGTTTGAGTGAGACGAGA | RT-qPCR |
| Pre-rRNA R | ACGTGCGCTCACCGAGAGCAG | RT-qPCR |
| rDNA Promoter F | GGTATATCTTTCGCTCCGAG | ChIP-qPCR, DRIP-qPCR |
| rDNA Promoter R | GACGACAGGTCGCCAGAGGA | ChIP-qPCR, DRIP-qPCR |
| 5'ETS-18S Junction F | GCCGCGCTCTACCTTAC | RT-qPCR |
| 5'ETS-18S Junction R | GCCGTGCGTACTTAGACAT | RT-qPCR |
| ss-Tag_hIGS-16 FS-A | CGAGGATCATGGTGGCGAATAAA CACACACACACCCCGTAGT | ss-RT-qPCR |
| ss-Tag_hIGS-16 FS-S | CGAGGATCATGGTGGCGAATAAG AAATGGGGCTTCGATACAT | ss-RT-qPCR |
| ss-Tag_hIGS-18 FS-A | CGAGGATCATGGTGGCGAATAAG TTGACGTACAGGGTGGACTG | ss-RT-qPCR |
| ss-Tag_hIGS-18 FS-S | CGAGGATCATGGTGGCGAATAAG GAAGTTGTCTTCACGCCTGA | ss-RT-qPCR |
| ss-Tag_hIGS-20 FS-A | CGAGGATCATGGTGGCGAATAAG TAGCCTTGGGCTTCTCTCC | ss-RT-qPCR |

| | | |
|---|---|---|
| ss-Tag_hIGS-20 FS-S | CGAGGATCATGGTGGCGAATAAA GTTTTCAGCCCCAACACAC | ss-RT-qPCR |
| ss-Tag_hIGS-22 FS-A | CGAGGATCATGGTGGCGAATAAC AGTGGCTCACGTCTGTCAT | ss-RT-qPCR |
| ss-Tag_hIGS-22 FS-S | CGAGGATCATGGTGGCGAATAAC GCCTGACTCCATTTCGTAT | ss-RT-qPCR |
| 7SK FS-A | TAATACGACTCACTATAGGGAGG ACCGGTCTTCGGTCAA | ss-RT-qPCR |
| 7SK FS-S | TAATACGACTCACTATAGGGTCAT TTGGATGTGTCTGCAGTCT | ss-RT-qPCR |
| ss-Tag | CGAGGATCATGGTGGCGAATAA | ss-RT-qPCR |
| 7SK-Tag | TAATACGACTCACTATAGGG | ss-RT-qPCR |
| 7SK RNA F | AGGACCGGTCTTCGGTCAA | ss-RT-qPCR, RT-qPCR |
| 7SK RNA R | TCATTTGGATGTGTCTGCAGTCT | ss-RT-qPCR, RT-qPCR |
| LINE 1 F | TGCGGAGAAATAGGAACACTTTT | ChIP-qPCR |
| LINE 1 R | TGAGGAATCGCCACACTGACT | ChIP-qPCR |
| RPLP1A Promoter F | TAAGAGGCTGCGTATAGGCG | ChIP-qPCR, |
| RPLP1A Promoter R | ATGAGGGCCGAGTAGATGCA | ChIP-qPCR, |
| RPLP1A RNA F | TCATTCTGCACGACGATGAGG | RT-qPCR |
| RPLP1A RNA R | TTGCAAACAAGCCAGGCCAA | RT-qPCR |
| PAF1 F | GCTTGTGGTCAAACATCGGG | RT-qPCR |
| PAF1 R | AATCAGCATCACTGAGCCCC | RT-qPCR |
| TBPL1 F | TGCCAGTTACGAACCTGAAC | RT-qPCR |
| TBPL1 R | GCCCTGTTACTGTGATACTTCC | RT-qPCR |

**Supplementary Table 3. List of sgRNAs used in this study.**

| sgRNA | Sequence |
|---|---|
| IGS 20-1 | TCTCCATTCGGAAGCTTGAC |
| IGS 20-2 | AAGGCTCTGTGCATACGAAT |
| IGS 20-3 | AATGATGAGACCCCGTCTGT |
| Non-targeting | AAAUGUGAGAUCAGAGUAAU |

**Supplementary Fig. 1. Validation of sucrose gradient-based nucleolar enrichment and sub-cellular fractionation. (a)** Representative bright field images of non-fractionated whole cell (WC) and nucleolar-enriched samples (NOL) using sucrose gradient. **(b)** Electrophoresis and Ponceau stain analysis of unfractionated whole cells (WC) compared to nucleoplasm/cytoplasm-enriched samples (N/C) and nucleolus-enriched samples (NOL). **(c)** Western blot-based confirmation of the enrichment of the nucleolar protein nucleophosmin (NPM) in NOL samples and the enrichment of the nuclear-cytoplasmic protein HSP70 in the WC and N/C samples. **(a-c)** Experiments were performed using Flp-In 293 T-REx$^{TM}$ cells, and images are representative of three biologically independent experiments.

**Supplementary Fig. 2. Validating the compBioID approach combining sucrose gradient-based nucleolar enrichment with mTurbo-POL2RI-based biotinylation of proteins. (a)** Western blot showing tetracycline(Tet)-inducible expression of Flag-mTurbo-POL2RI or Flag-mTurbo in whole cells. **(b)** Western blots confirming mTurbo-dependent protein biotinylation in whole-cell samples from cells expressing Tet-inducible Flag-mTurbo-POL2RI or control Flag-mTurbo. **(c)** Western blot showing the differential biotinylation of proteins in non-fractionated whole-cell samples (WC) and nucleolar-enriched samples (NOL) following the Tet-dependent induction of Flag-mTurbo-POL2RI. **(a-c)** Experiments were performed using Flp-In 293 T-REx$^{TM}$ cells and images are representative of two biologically independent experiments.

**Supplementary Fig. 3. Additional characterization of compBioID-detected proximity interaction candidates of nucleolar Pol II. (a-c)** ChIP assays assessing the enrichment of TBPL1, PAF1, MED26 or DR1 at indicated genomic sites. Experiments were performed using HEK293T cells, data are shown as mean±s.d. with $n$ = 3 biologically independent experiments and statistical significance was assessed using multiple unpaired $t$-tests. **(d)** Portion of the IGS DNA sequence showing antisense TCT motifs around the IGS 22 and 32 sites. The positions of primers used for ChIP-based assays in this study are shown.

**Supplementary Fig. 4. Controls related to the characterization of the role of TBPL1 at the IGS. (a)** RT-qPCR analysis showing the impact of TBPL1 knockdown on average IGS ncRNA expression. **(b)** ChIP showing the relative change in the enrichment of Pol II at the ribosomal protein gene *RPLP1A*. **(c)** Immunoblots showing the impact of TBPL1 knockdown on Pol II levels. Vinculin served as loading control. **(d)** Confirmation of the sensitivity of DRIP signals across the IGS to *in vitro* RNaseH1 treatment in cells with siCTRL and siTBPL1. **(e)** Detection of R-loops across the IGS using catalytically inactive RNaseH1(DN) in the presence or absence of *in vitro* RNaseH1 treatment. **(f)** Impact of TBPL1 knockdown on R-loops as detected using RNaseH1(DN). **(g)** RT-qPCR-based analysis of the impact of TBPL1 knockdown on nascent RNA expression from the *RPLP1A* gene. **(h)** Immunoblots showing the impact of TBPL1 knockdown on Pol I levels. Vinculin served as loading control. **(i)** ChIP-re-ChIP shows the co-enrichment of TBPL1 with Pol I at its rRNA gene promoter. **(j)** Impact of TBPL1 knockdown on Pol I enrichment at the rRNA gene promoter. **(k)** Portion of the IGS DNA sequence showing the sense TCT motif proximal to the IGS16 site. Primer locations used for ChIP-based assays in this study are also shown. **(a-j)** HEK293T cells were used; data are shown as mean±s.d.; $n$ = 3 biologically independent experiments; unpaired $t$-test (a, b, g, i, j) or multiple unpaired $t$-tests (e, f) were used; images (c, h) are representative of three independent experiments.

**Supplementary Fig. 5. Controls related to the characterization of the role of PAF1 at the IGS. (a)** RT-qPCR analysis showing the impact of PAF1 knockdown on average IGS ncRNA expression. **(b)** Immunoblots showing the impact of PAF1 knockdown on Pol II levels. Vinculin served as loading control. **(c)** Confirmation of the sensitivity of DRIP signals across the IGS to *in vitro* RNaseH1 treatment in cells with siCTRL and siPAF1. (**d**) Impact of PAF1 knockdown on R-loops as detected using RNaseH1(DN). **(e)** Immunoblots showing the impact of PAF1 knockdown on Pol I levels. Vinculin served as loading control. **(f, g)** ChIP analysis confirmed the enrichment of RED (f) or dRED (g) at the IGS20 site using sgRNAs targeting that site (sg20). Non-targeting sgRNA (sgNT) served as control. **(h, i)** Use of the LasR system with the dRED fusion protein and sg20 represses did not repress R-loops at the IGS20 site in DRIP (h) or increases IGS ncRNA levels at the IGS in RT-qPCR (i). **(a-i)** HEK293T cells (a-e) or HEK293 TREx$^{TM}$ cells (f-i) were used; data are shown as mean±s.d.; *n* = 3 (a-f, i) or *n* = 2 (g, h) biologically independent replicates; unpaired *t*-test (a, d, f) or ordinary two-way ANOVA with uncorrected Fisher's LSD test (c, i) were used; images (b, e) are representative of three independent experiments.

**Supplementary Fig. 6. Uncropped blots.** Shown are uncropped blots together with a dashed box outlining the cropped area and the molecular weight ladder in kDa.

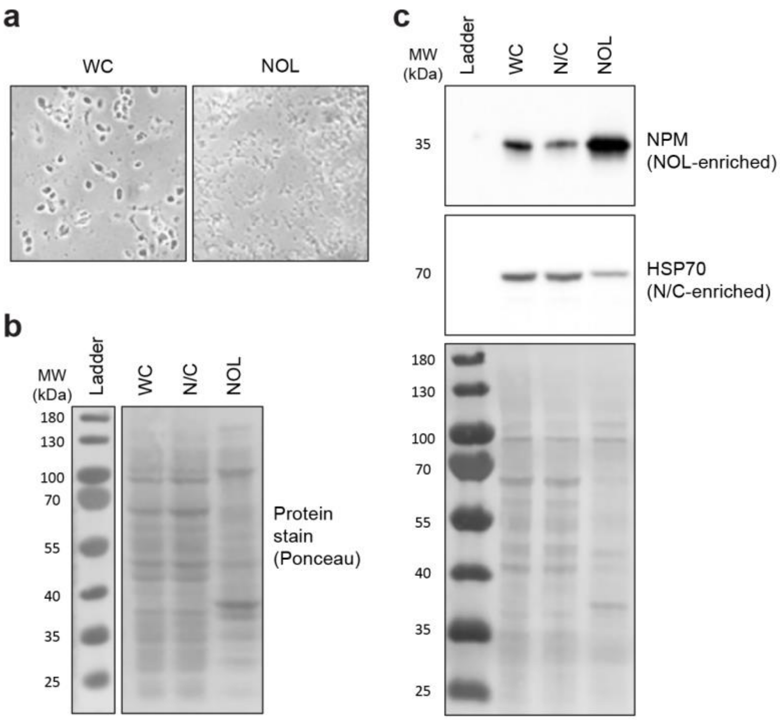


**Supplementary Fig. 1**

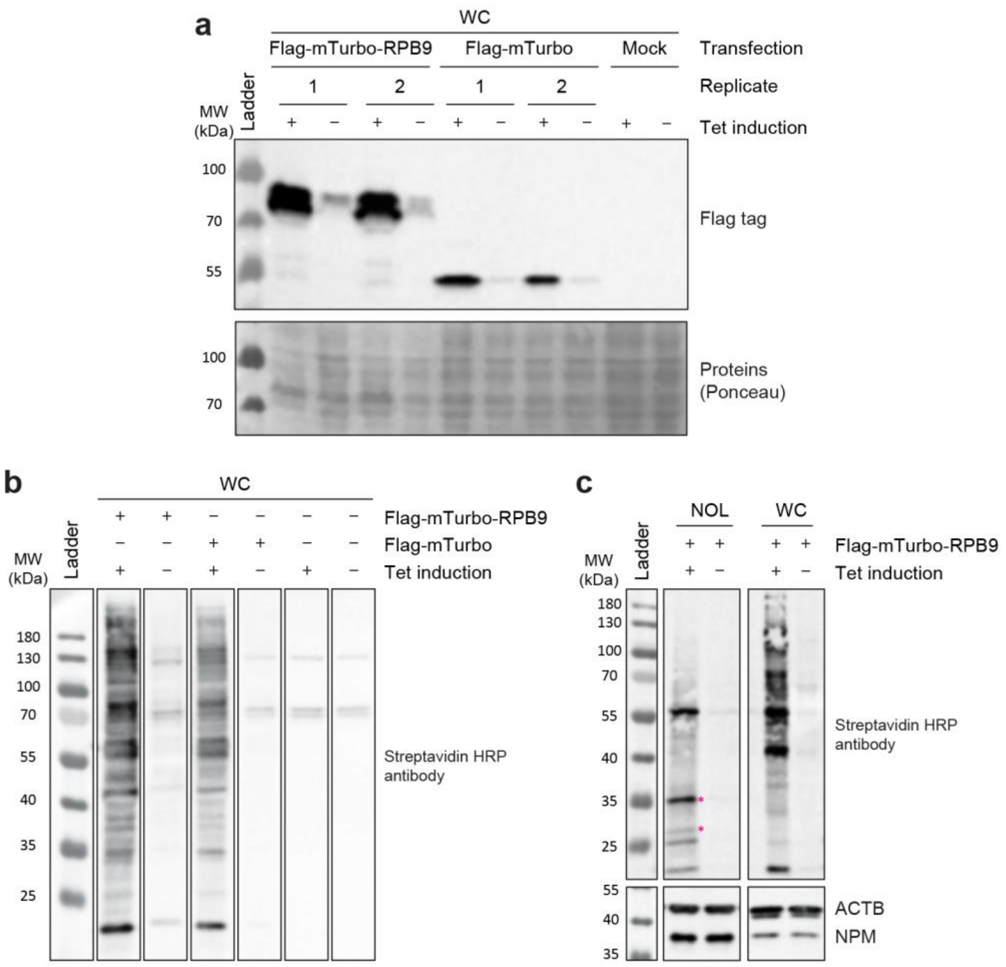


### Supplementary Fig. 2

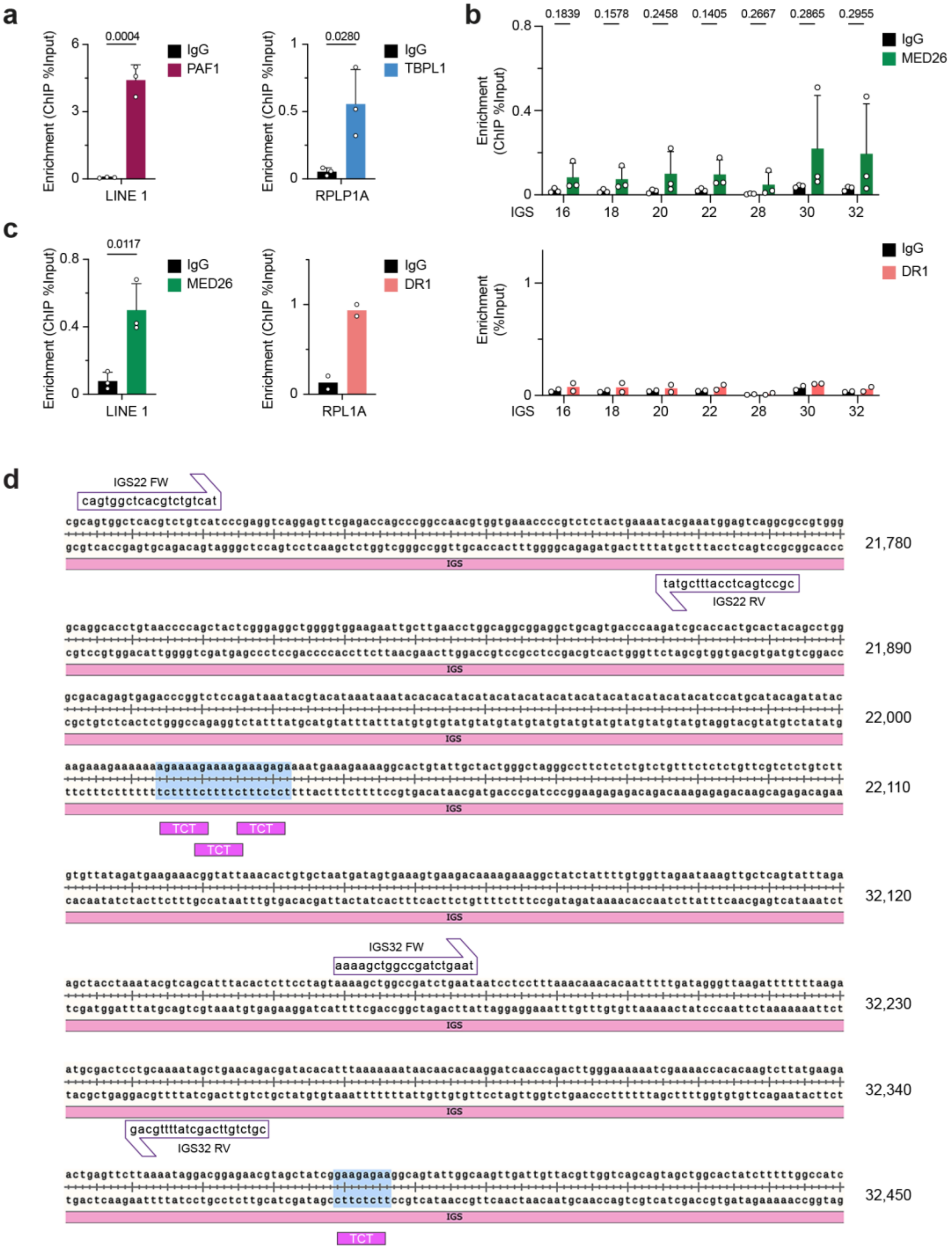


### Supplementary Fig. 3

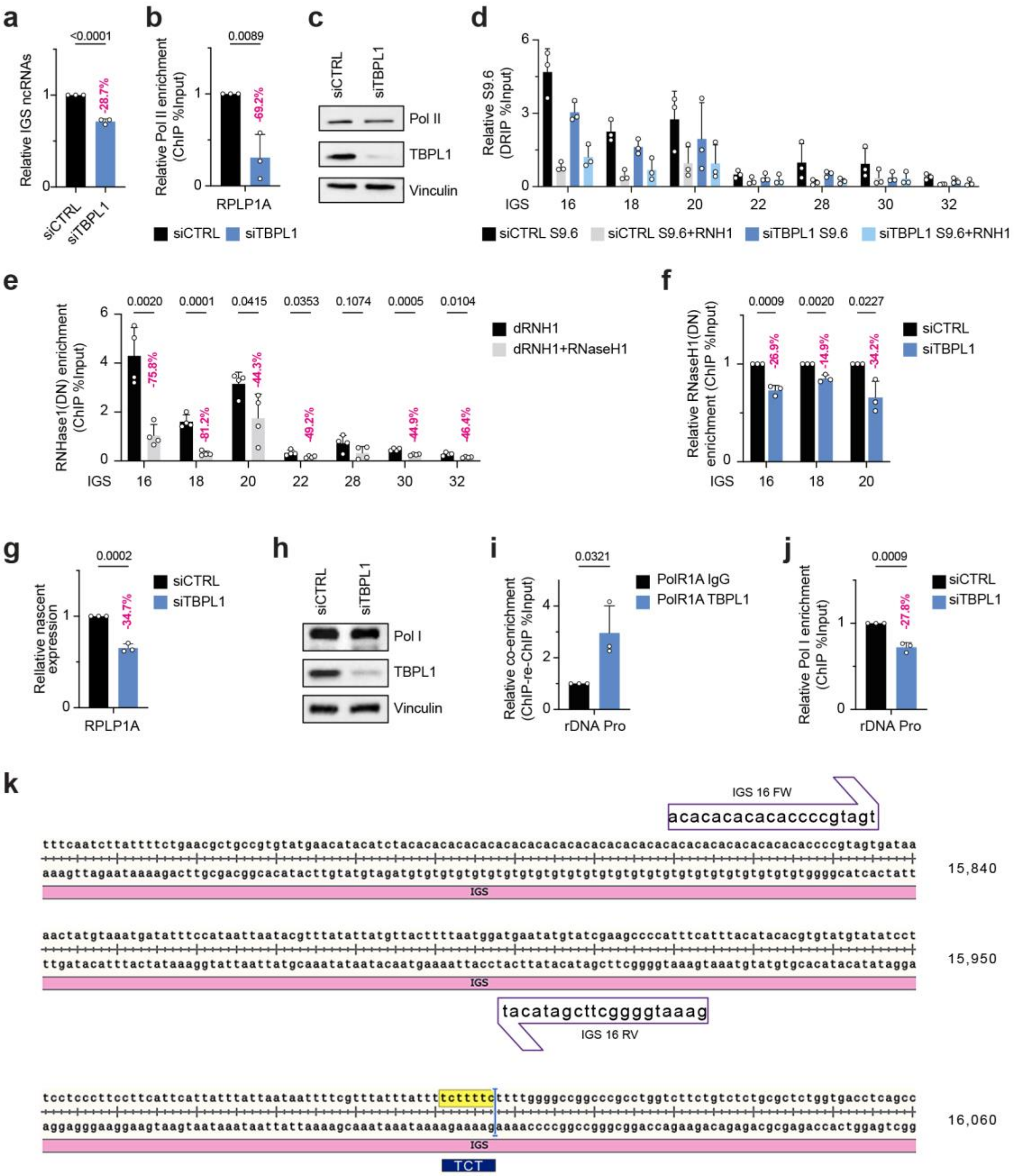


**Supplementary Fig. 4**

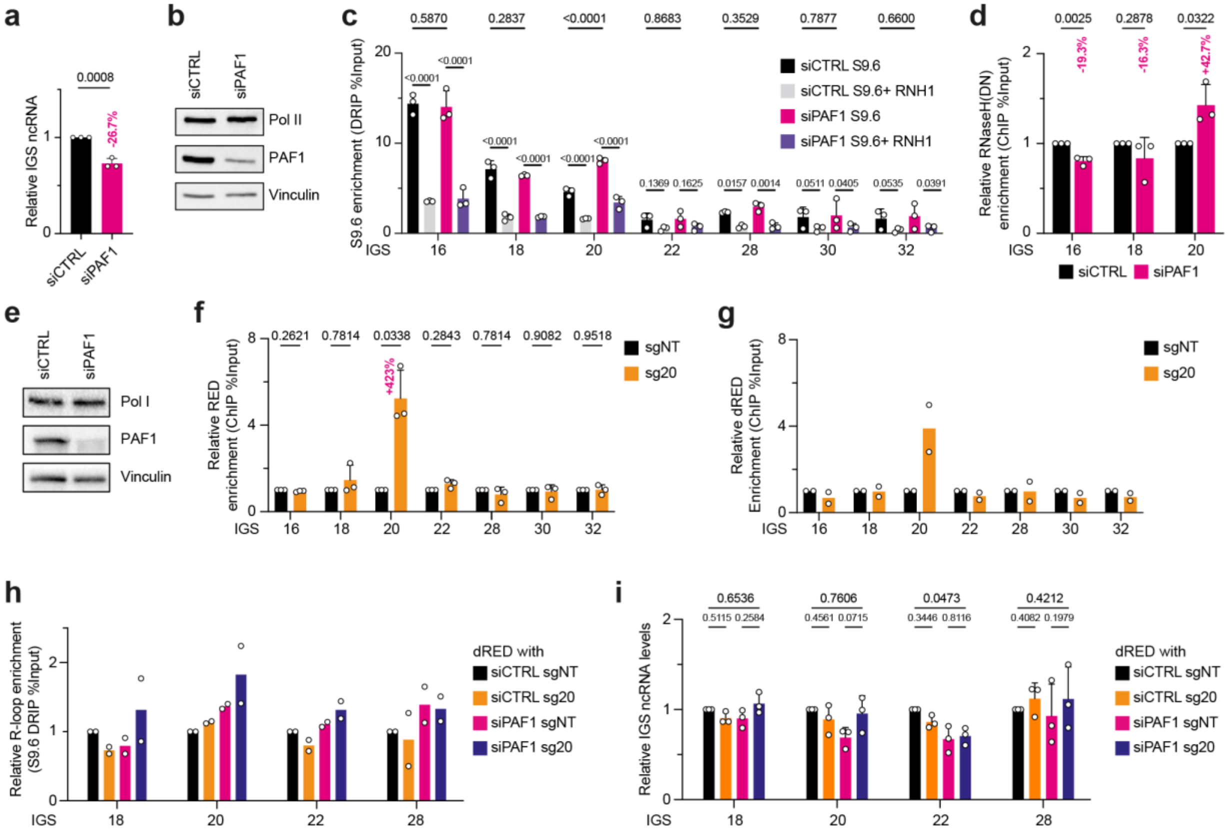


Supplementary Fig. 5

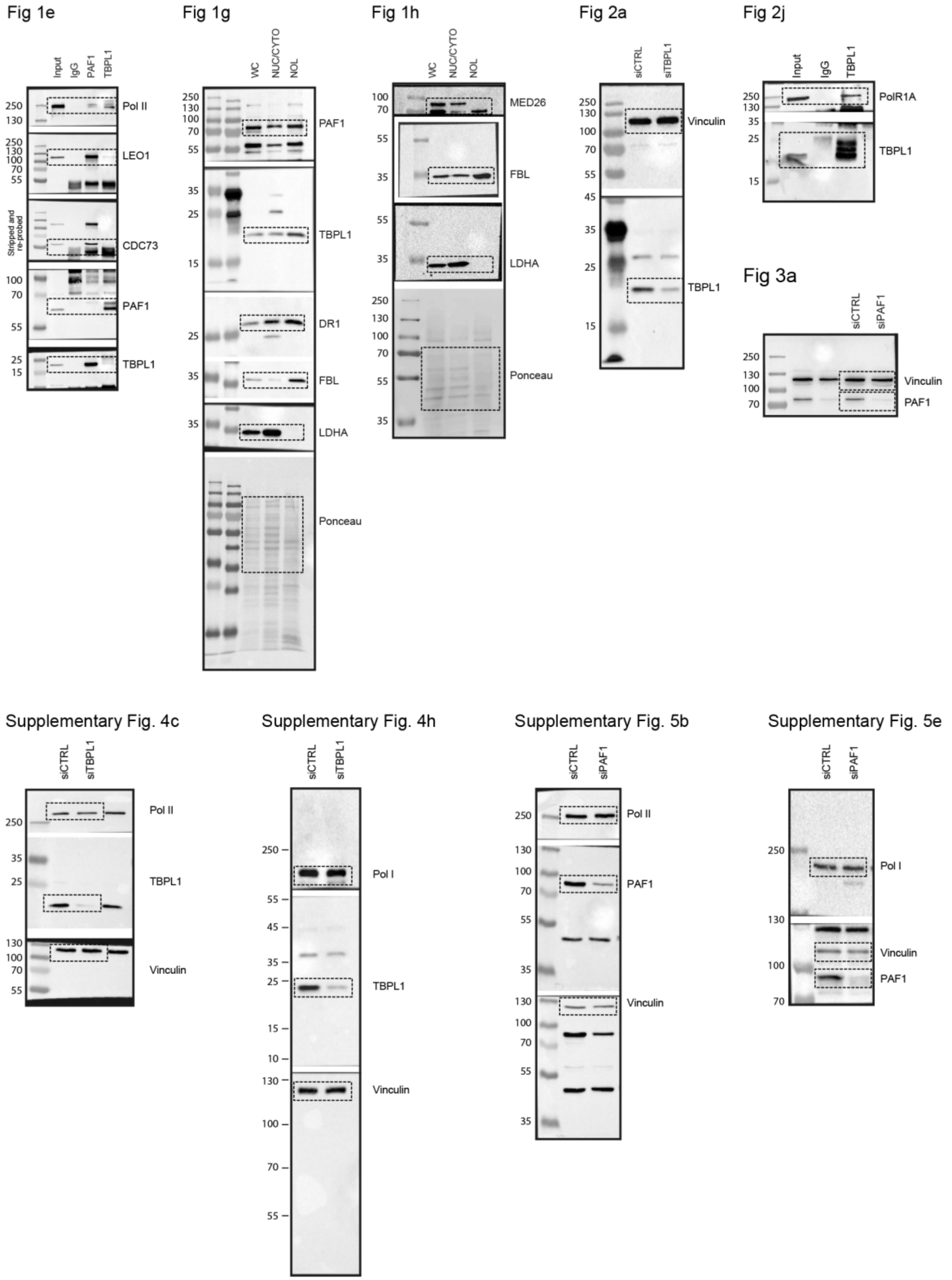


**Supplementary Fig. 6**
